# Host genetics and the Skin Microbiome Independently Predict Parasite Resistance

**DOI:** 10.1101/2024.10.19.619196

**Authors:** Rachael D. Kramp, Mary J. Janecka, Nadine Tardent, Jukka Jokela, Kevin D. Kohl, Jessica F. Stephenson

## Abstract

Host responses to parasite infection involve several interacting systems. Host genetics determine much of the response, but it is increasingly clear that the host-associated microbiome also plays a role. Genetically determined systems and the microbiome can also interact; for example, the microbiome can modulate the immune response, and vice versa. However, it remains unclear how such interactions between the host immune system and the microbiome may influence the host’s overall response to parasites. To investigate how host genetics and the microbiome interact to shape responses to parasites, we imposed truncation selection on Trinidadian guppies (*Poecilia reticulata*) for low and high resistance to the specialist ectoparasite *Gyrodactylus turnbulli*. After 3-6 generations of breeding without parasites, we sampled the skin-associated microbiome and infected fish from each line. We applied Dirichlet Multinomial Modeling (DMM) machine-learning to identify bacterial community types across lines and evaluated how selection line and community type explained variations in infection severity. Our findings showed that among females, the resistant line had significantly lower infection severity, while the susceptible line had higher infection severity. Among males, only the susceptible line experienced higher infection severity compared to the other lines. Line did not explain skin microbial diversity, structure or composition. Our DMM analysis revealed three distinct bacterial community types, independent of artificial selection lines, which explained just as much variation in infection load as selection line. Overall, we found that the microbiome and host genetics independently predict infection severity, highlighting the microbiome’s active role in host-parasite interactions.

## 1. Introduction

Hosts use multiple strategies to defend themselves against parasites and pathogens (parasites hereafter). Host genetic background is commonly credited with much of the defense against parasites through the immune system, behavior, or other genetic/protein alterations, e.g., thalassemia (Labro, 2022). Recent research has highlighted that the host-associated microbial communities, or ‘microbiome’, may also influence parasite defense indirectly through the host immune system or directly via niche occupation or secreting anti-parasitic compounds (Bernardo-Cravo et al., 2020), and may also exhibit heritable components. Indeed, the host-associated microbiome can dramatically impact host survival and defense against parasites. However, the question of whether the microbiome plays an active role in host-parasite infection or merely reflects the host’s health/immune status is commonly posed (Trevelline et al., 2020). While there is abundant evidence that the microbial community can have a direct or indirect role in host-parasite interactions (Becker et al., 2009; Nelson & May, 2020), there is also evidence that the microbiome is merely a bystander in some diseases and a reflection of the host’s genetics, diet, and general health (Ni et al., 2017).

Host genetics and the microbiome interact throughout host development, with complex feedbacks complicating the separation of their roles in traits like parasite defense. On one hand, hosts exert control over their symbionts, shaping their associated microbiomes, via immune responses, barrier function, and physiological regulation (Wilde et al., 2024). On the other hand, the host immune system relies on early microbial colonization for proper development and ultimately shapes immune cell composition (Amenyogbe et al., 2017). The host’s genetic makeup therefore influences microbiome composition, and may lead to the selection of specific microbial communities that can be inherited across generations (Bordenstein & Theis, 2015). However, distinguishing host genetic effects on the microbiome from factors such as priority effects, drift, and other stochastic elements, is an ongoing challenge (Debray et al., 2022), and the influence of external factors such as environment and temporal changes in shaping host-associated microbiomes must also be carefully considered. Direct manipulation of the host genetics in traits like parasite resistance, where the microbiome likely plays a role, thus provides a useful tool in disentangling host genetic effects from microbial ones.

Both the host-associated microbiome and the immune system commonly differ between host sexes (Markle & Fish, 2014). These differences are likely due to other correlated differences, for example, high testosterone levels in the blood both suppress the immune system and significantly alter gut microbial communities (Santos-Marcos et al., 2023), and both the immune system and microbiome change during pregnancy and menstruation (Chen et al., 2023; Gorczyca et al., 2022). Given that host sexes across taxa also differ in their response to parasites (Morrow, 2015), it is essential to test for sex-specific interactions between genetic resistance factors, the microbiome, and parasites.

Trinidadian guppies (*Poecilia reticulata*) and their specialist ectoparasite *Gyrodactylus turnbulli* provide a rich context to study host-parasite dynamics and interactions with the microbiome. *Gyrodactylus* spp. feed and reproduce (both sexually and asexually, generation time 24hrs) on host skin. Guppies show a large variation in parasite resistance across populations, sexes, and ecological contexts (Dargent et al., 2016; Phillips et al., 2021; Stephenson, van Oosterhout, & Cable, 2015; Van Oosterhout et al., 2003). Guppy resistance is at least partly genetically determined and heritable (Dargent et al., 2016; Phillips et al., 2018, 2021; Van Oosterhout et al., 2003; Weiler et al., 2025). Innate and acquired immunity are important for *G. turnbulli* resistance, however, the precise mechanisms involved in guppy parasite resistance remain unclear (Cable & Van Oosterhout, 2007; Mohammed et al., 2016). Research into genetically based resistance to *G. turnbulli* has largely focused on the major histocompatibility complex (MHC) (Fraser et al., 2010; Fraser & Neff, 2009; Phillips et al., 2018). Notably, skin bacterial alpha diversity correlates with MHC heterozygosity in frogs (Cortazar-Chinarro et al., 2024), and probiotic supplementation results in significant upregulation of MHC genes in fish (Tanpichai et al., 2025). Similar associations and covariation between host genetics and the skin microbiome of fish would sets up a clear avenue for the skin microbiome and host genetics to interactively shape parasite resistance phenotypes in our system.

Here, we tested how host genetics and host-associated microbiomes affect the interaction between guppies and *G. turnbulli*. We imposed truncation selection for host resistance and susceptibility to the parasite, quantified as the mean number of parasites over infection. While we did not specifically select for tissue-specific tolerance (tolerance hereafter), or the ability to minimize the fitness losses caused by a given parasite load (Stephenson & Adelman, 2022), we did examine the levels of tolerance within our lines. Previous literature has demonstrated that tolerance is an essential factor to consider in guppy response to *Gyrodactylus* parasites (Jog et al., 2022; Stephenson, van Oosterhout, & Cable, 2015; Tadiri et al., 2021). However, whether tolerance of infection is heritable has rarely been explored.

We found that host genetics and the microbiome significantly and independently predicted the severity of parasite infection. Parasite resistance and tolerance showed sex-specific responses to selection. These results suggest that the microbiome independently influences host-parasite interactions rather than merely reflecting the host’s genetics, highlighting the complex interplay among host genetics, the microbiome, and sex-specific responses to infections.

## 2. Materials and methods

### 2.1 Data and sample collection

#### 2.1.1 Truncation selection to establish guppy lines that differ in genetically-based resistance to Gyrodactylus turnbulli

We used laboratory-bred descendants of wild guppies from a high-predation population on the Caura River, Trinidad (UTM: 20 P 679527.7 m E, 1180376.4 m N; elevation 112 m). Wild guppies (n ∼600) were transported to Cardiff University in June 2012 (Cefas APB authorization number CW054-D-187A) and treated prophylactically with Binox® (Nitrofurazone). The site was previously confirmed *Gyrodactylus* spp.-free by previous surveys, and no *Gyrodactylus* spp. were found in a subset of 80 fish examined. We collected fin clips from 66 parental-generation fish for microsatellite genotyping. Fish were housed in 70 L mixed-sex tanks under a 12h light:12h dark cycle at 24±1°C and fed daily with Aquarian® tropical fish flakes, supplemented with Artemia and bloodworm. Fry were transferred to juvenile tanks soon after birth, checked for ectoparasitic infection, and reared until sex determination (6-8 weeks). They were then kept in sex-specific tanks for at least another month before use, ensuring all fish were sexually mature and unmated. *Gyrodactylus turnbulli* were from an established isogenic culture in the lab (*Gt3*), maintained on commercially sourced guppies (‘culture fish’).

We experimentally infected 100 male and 100 female F1 guppies to evaluate their resistance and select founders for our ‘resistant’ and ‘susceptible’ lines. To initiate infections, a heavily infected culture fish was euthanized with MS222 (0.02% tricaine methanesulfonate) and placed next to an anesthetized experimental fish until two *G. turnbulli* parasites transmitted, as observed under a dissecting scope. Experimental fish were housed individually in 1 L tanks, and parasites were counted every other day on anesthetized fish. After nine days, all fish were treated with Levamisole and screened under anesthetic several times to confirm parasite clearance. Fin clips were collected for genotyping. The 30 males and 30 females with the highest (lowest) mean number of worms founded the ‘susceptible’ (‘resistant’) line, and randomly selected uninfected F1 fish (30 of each sex) founded the control line (*Figure S1*). Each line was allowed to breed freely in separate 120 L tanks. Tanks were monitored for fry, which were moved to F2 rearing tanks. The F2 generation was split into two subpopulations (named ‘A’ and ‘B’ within each of the susceptible, resistant, and control lines), maintaining 60 fish per tank. This population size was kept consistent across the six subpopulations, with new tanks set up for each subsequent generation as above. During preliminary analysis, we confirmed that these subpopulations did not differ in either infection severity or parasite tolerance (see Supporting Information) and therefore combined them for the subsequent analyses described below.

After roughly 3 generations post-selection, the fish moved with one of the authors to a new institution, at which the husbandry conditions were different. Here, guppies were housed in mixed-sex groups at densities of 1-2 fish per L in 4.5 L tanks in a recirculating system at 24±1°C, on a 12 h light: 12 h dark lighting schedule (overhead fluorescent lighting) and fed daily on commercial flake food supplemented with *Artemia*. Each subpopulation was therefore housed across at least 10 separate tanks, again with multiple new tanks set up for each subsequent generation. The fish were held and bred under these conditions from August 2015 until their experimental infections in August-November 2017.

#### 2.1.2 A note on replication in our study design

As described above, we imposed one round of truncation selection for resistance and susceptibility, on one group of F1 fish. Typically, selection experiments using guppies to characterize the genes underlying phenotypic variation (e.g., (van Wijk et al., 2013)), or evolutionary change in correlated traits (e.g., (Bartuseviciute et al., 2022; Kotrschal et al., 2013)) replicate this process three times because the scale of inference is at the level of the selection event: to draw conclusions about how underlying genes or correlated traits respond to selection, the selection event should be replicated (Kawecki et al., 2012).

By contrast, we do not identify genes, or focus on correlated evolutionary trait changes: instead, we use truncation selection to maximize the variation in response to infection in our focal population, and to compare the role of host genetics and associated microbiomes in infection phenotypes. That resistance to *Gyrodactylus* spp. is heritable is not our focus, and has already been established several times (Dargent et al., 2013; Madhavi & Anderson, 1985; Phillips et al., 2018, 2021; Weiler et al., 2025). To ensure that the difference between our lines was due to selection, and thus underlying genetic differences, we kept consistent population sizes across generations, split each line into two subpopulations, housed fish across several replicate tanks per line, bred the fish for at least 3 generations in the absence of parasites, and used genotyping to eliminate the role of inbreeding differences between them. With our design, therefore, we can rule out potential confounding effects (from tanks, stochastic differences post selection, inbreeding differences, and maternal effects).

Our selection process thus produced individual guppies that differed in genetically-based resistance to our focal parasite; here we show that the microbiomes associated with these individuals contributed independently to their infection phenotype. Our main conclusion, that the host-associated microbiome predicts infection phenotype independently of and just as strongly as host genetics, is therefore robustly supported by our study design.

#### 2.1.3 Experimental infection and microbiome sampling protocol

Before infection began, we collected samples from 51 fish split between the infection and sham-infection treatment groups: control (infected n=10, sham n=5), Resistant (infected n=12, sham n=6), and Susceptible (infected n=9, sham n=8). We used the methods previously published to collect and inventory the microbial communities on the fish skin.

We infected 88 guppies 3-7 generations post truncation selection with *G. turnbulli* in five batches, following the same procedure as for the F1 generation, described above. 14 guppies from each line were ‘sham’ infected and were used in microbiome analysis. *G. turnbulli for this infection* were collected from a local pet shop in June 2017. An isogenic strain was initiated by infecting a parasite-naïve ornamental guppy with a single parasite individual. This strain was subsequently maintained on parasite-naïve culture fish, and identified to species level using molecular methods.

### 2.2 What is the magnitude of host genetic contribution to guppy resistance to Gyrodactylus turnbulli?

#### 2.2.1 Statistical analysis

We used R v.4.3.2 (R Core Team, 2020) for these analyses and provide our code, output, and validation in the Supporting Information (SI). We used ggplot2 (Wickham, 2016) to plot all the figures. For all models, we assessed collinearity among predictor variables using the pairs function (Zuur et al., 2009), validated model fits using the DHARMa package (Hartig, 2022), and used Type II Wald chi-square tests to test for significance using the *Anova()* function from the ‘car’ package (Fox & Weisberg, 2019).

We quantified infection resistance/susceptibility as the area under the curve of parasite number over time, or “infection severity.” To test response to selection, we used infection severity as the response variable in a generalized linear mixed model (GLMM) in the glmmTMB package, v. 1.1.2.3 (Brooks et al., 2017) (Gamma error family and logit link function). As fixed effects, we included selection line (resistant n=30, control n=28, susceptible n=29), fish sex, length (pre-infection, using residuals from a regression on sex to control for sex differences), body condition (scaled mass index; (Peig & Green, 2009)), generation, number of parasites establishing the infection (dose), and trial date (Julian date). Length and body condition were calculated with entire population of fish including sham infected fish (resistant n=14, control n=14, susceptible n=14), to obtain a more accurate distribution. We also included the interaction between sex and line.

We quantified tolerance as the host’s relative per-parasite percentage mass change during infection, calculated by dividing the percentage mass change from day 0 to day 20 by the maximum number of worms on the fish, relative to the highest observed percentage change. To assess the impact of selection on tolerance, we used tolerance as a response variable in a GLMM, including the fixed effects of line, sex, length, dose, and Julian date. We excluded scaled mass index due to its overlap with mass change. Furthermore, we did not include infection severity due to high collinearity with other fixed effects within the model (line & dose). Additionally, infection severity within the original model caused model fit issues due to the non-linear relationship between infection severity and ppmass.

In a different model, we tested for a tradeoff between resistance and tolerance with a quadratic polynomial regression model to account for non-linear effects. We used tolerance as the response variable and log-transformed infection severity (our resistance metric), sex, their interaction, length, and generation as fixed effects.

#### 2.2.2 Calculating heritability

We calculated narrow-sense heritability (assuming that the response to selection is primarily due to additive genetic variance) of resistance (using mean worm – the resistance metric we used during truncation selection) and tolerance following the breeder’s equation (Lush, 1937). For mean worm, we used the average number of parasites between 1-9 days of infection, centered to a mean of zero to account for the differences in worm growth between the infections of the F1 and subsequent generations. Tolerance was centered identically.

P_b_: Average starting phenotype (mean phenotype of the population before selection)

P_s_: Average selected phenotype (mean phenotype of the selected parents)

P_r_: Average response phenotype (mean phenotype of the offspring generation)

h^2^: Narrow-sense heritability

h^2^= P_r_-P_b_P_s_-P_b_

#### 2.2.3 Do differences in inbreeding contribute to line differences?

We evaluated the genetic diversity and variance in inbreeding within the wild-caught populations, F1, and F3-F6 generations, to determine if the observed difference between the resistant and susceptible lines could be attributed to differences in inbreeding. We extracted DNA from fin clips with the HotSHOT method: using 50 µl of alkaline lysis reagent and neutralizing solution followed by incubation at 95 °C for 30 minutes (Truett et al., 2000). We used two multiplex assays to amplify eight previously designed microsatellite markers (Becher et al., 2002; Nater et al., 2008; Paterson et al., 2005; van Oosterhout et al., 2006; Watanabe et al., 2005) at the Genetic Diversity Centre (GDC), ETH Zurich. Samples were genotyped on an ABI Genetic Analyzer 3730 (Applied Biosystems, USA), scored using GeneMarker (Version 2.6.4; Softgenetics), and binned using the package MSatAllele (Alberto, 2013) in R (R Core Team, 2020). See the supporting information for more details.

We tested allelic richness (*A_R_*), gene diversity (*H_S_*), *F_IS,_* and identity disequilibrium (g2) among our artificial selection lines to evaluate whether the differences in phenotype were attributable to variation in inbreeding (Smallbone et al., 2016). Allelic richness is the first population genetic metric to reflect bottleneck effects, which decrease the effective population size and facilitate reductions in heterozygosity due to inbreeding (Nei et al., 1975). Significant differences in allelic richness among populations are indicative of variation in increases in the probability that inbreeding may occur. Gene diversity measures the mean heterozygosity per individual per population, under inbreeding conditions homozygosity increases but gene diversity is not immediately lost (Charlesworth, 2003). Positive values in *F_IS_* can be generated by inbreeding but may also be artificially elevated by technical artifacts such as large allele drop out or null alleles, which can artificially inflate a deficiency of heterozygotes (De Meeûs, 2018; Nei et al., 1975). We therefore also estimated identity disequilibrium (*g*_2_), the correlation of heterozygosity across loci resulting from inbreeding, which is robust in the presence of null alleles (David et al., 2007). Concurrence among population genetic metrics consistent with patterns inbreeding are needed to determine if phenotypic differences among populations are attributable to inbreeding. Allelic richness (*A_R_*) (rarefied to the lowest population sample size), gene diversity (*H_S_*) and *F_IS_*, was calculated in FSTAT 2.39 (Goudet, 01 2001). Significance levels for *F_IS_* were assessed in GENEPOP (web version 4.2;(Raymond & Rousset, 1995) with 5000 iterations followed by a Bonferroni correction for multiple comparisons. Tests for statistically significant differences allelic richness (*A_R_*), gene diversity (*H_S_*) and *F_IS_* among groups were calculated in FSTAT with 2000 permutations. Identity disequilibrium (*g*^2^) is the difference between joint identity by descent and the product of the separate probabilities of identity by decent for two loci. *g^2^*is estimated with microsatellite markers as the excess of double heterozygous genotypes at all pairs of loci relative to a random distribution of heterozygous genotypes among individuals at each locus (Cockerham & Weir, 1973; David et al., 2007). *g_2_* values significantly greater than 0 are indicative of identity disequilibrium (ID). ID was calculated in inbreedR (Stoffel et al., 2016).

### 2.3 Does the host-associated microbiome contribute to guppy resistance to Gyrodactylus infection?

#### 2.3.1 Microbial alpha and beta diversity

Raw sequence data were processed using QIIME2 version 2019.7 (Bolyen et al., 2019). We rarefied the resulting ASV table to 3,654 sequences per sample which resulted in excluding one sample and kit controls, and used the DADA2 pipeline to calculate the unweighted and weighted UniFrac distances (Lozupone & Knight, 2005) pairwise between all samples to create distance matrices. We used a PERMANOVA from the adonis package in R (Anderson, 2001) to test for significant differences in microbial community across fish line, sex, length, and body condition. We used QIIME2 to calculate the number observed number of ASVs, Shannon’s diversity index, and Faith’s phylogenetic diversity, and compare the mean values of these metrics across line and sex using Pairwise Kruskal-Wallis tests.

#### 2.3.2 Bacterial community typing

We used Dirichlet multinomial mixture models (DMM) (Holmes et al., 2012) in the Dirichlet Multinomial v1.16.0 package (Morgan, 2023) to assign all samples (n=50) to three different bacterial community types based on the relative frequency of taxa. We compared the Laplace approximation of the negative log models to identify the number of Dirichlet components that resulted in the best model fit (see SI).

#### 2.3.3 Does bacterial community type predict the severity of Gyrodactylus infection?

To test if the bacterial community type significantly predicted infection severity, we used bacterial community type as a fixed effect in the GLMM model used for the selection line resistance analysis described in section 2.1.2, with the data subset to just swabbed fish.

### 2.4 What is the relative importance of host genetics and the microbiome to guppy resistance to Gyrodactylus infection?

To compare how well our GLMM models with a) bacterial community type only, b) artificial selection line only, and c) both factors explained variation in infection severity, we used the *anova*() function. All three models shared the same fixed effects, differing only in whether line or bacterial community type or both were included as a fixed effect. Models involving skin microbiome and infection were run on the reduced dataset using only the infected and swabbed fish (n=31).

## 3. Results

### 3.1 Host genetics contribute to guppy resistance to, and tolerance of, Gyrodactylus turnbulli infection

#### 3.1.1 Resistance responded to selection

Selection line was a significant predictor of infection severity (Figure 1; χ^2^(2) =37.74, p=6.4e-09) Additionally, an interaction between selection line and sex was observed (χ^2^(2) =13.51, p=0.0012). Females had significantly higher infections than males (χ^2^(1) =24.29, p=8.3e-07) and high starting worm ‘dose’ resulted in higher infection severity (χ^2^(1) =5.11, p=0.024). Post hoc tests to evaluate whether selection line significantly explained infection severity among each sex found that line was still significant in both sexes (female; χ^2^(2) =24.33, p=5.2e-06 & male; χ^2^(2) =17.57, p=0.00015). Post hoc Tukey tests also revealed that the female guppies from the susceptible line experienced significantly higher (Cohen’s d=3.49, p<0.001), and from the resistant line experienced significantly lower infection severity (Cohen’s d=2.74, p=0.016) than the control. Male guppies from the susceptible line experienced significantly higher infection severity than control line males (Cohen’s d=4.19, p<0.001), but resistant line males did not differ significantly from control line males (Cohen’s d=1.39, p=0.15). The heritability of the average number of parasites throughout infection was h^2^=0.79 in the susceptible line, and h^2^=0.41 in the resistant line.

**Figure 1.**
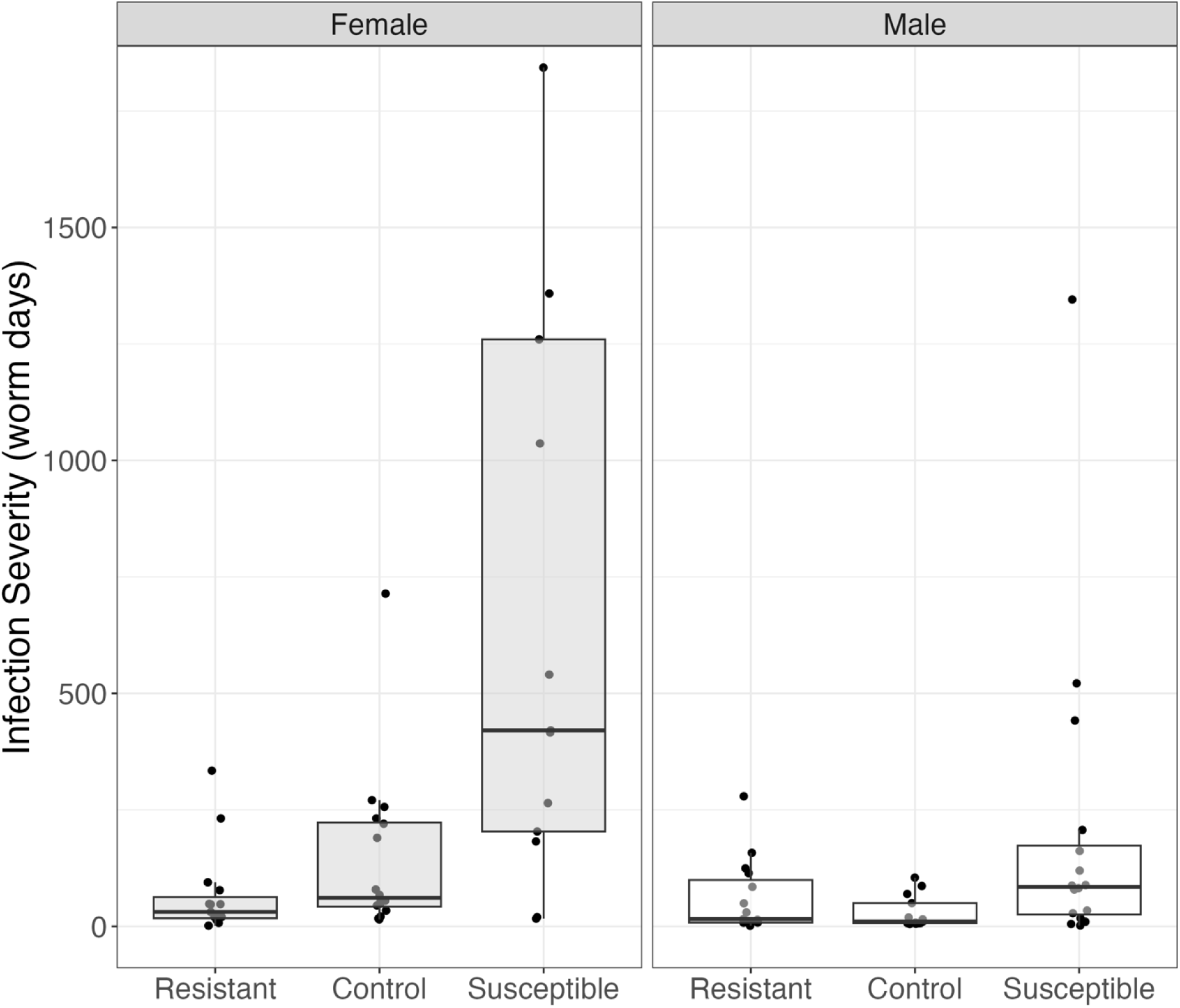
Artificial selection lines significantly predicted G. turnbulli infection severity in Trinidadian guppies. Infection severity (worm days) reflects the area under the infection load curve. Female guppies (n=44) in susceptible and resistant lines differed from the control line. Male guppies (n=43) only showed differences in the susceptible line. Box plots illustrate median, interquartile range, and values within 1.5× the interquartile range of infection severity.

#### 3.1.2 Tolerance responded to selection and trades off with resistance

The selection lines significantly differed in relative tolerance (Figure 2; χ^2^(2)=14.56, p=0.00069) and significantly interacted with sex (χ^2^(2)=10.28, p=0.00586). Additionally, larger fish were more tolerant (χ^2^(1)=5.94, p=0.014). A post hoc test found that selection line was only significant in female guppies (female; χ^2^(2)=23.81, p=6.7e-06), and smaller females were less tolerant (χ^2^(2) =6.96, p=0.0083). Post hoc Tukey tests revealed that resistant line females were significantly less tolerant than control and susceptible line females (control; Cohen’s d=7.04, p=0.0003; susceptible; Cohen’s d=8.54, p<0.00016). However, no fixed effect explained variation in tolerance among males and we did not find that the bacterial community types significantly predicted tolerance. The heritability of tolerance was h^2^=0.52 in the susceptible line and h^2^=0.24 in the resistant line.

**Figure 2.**
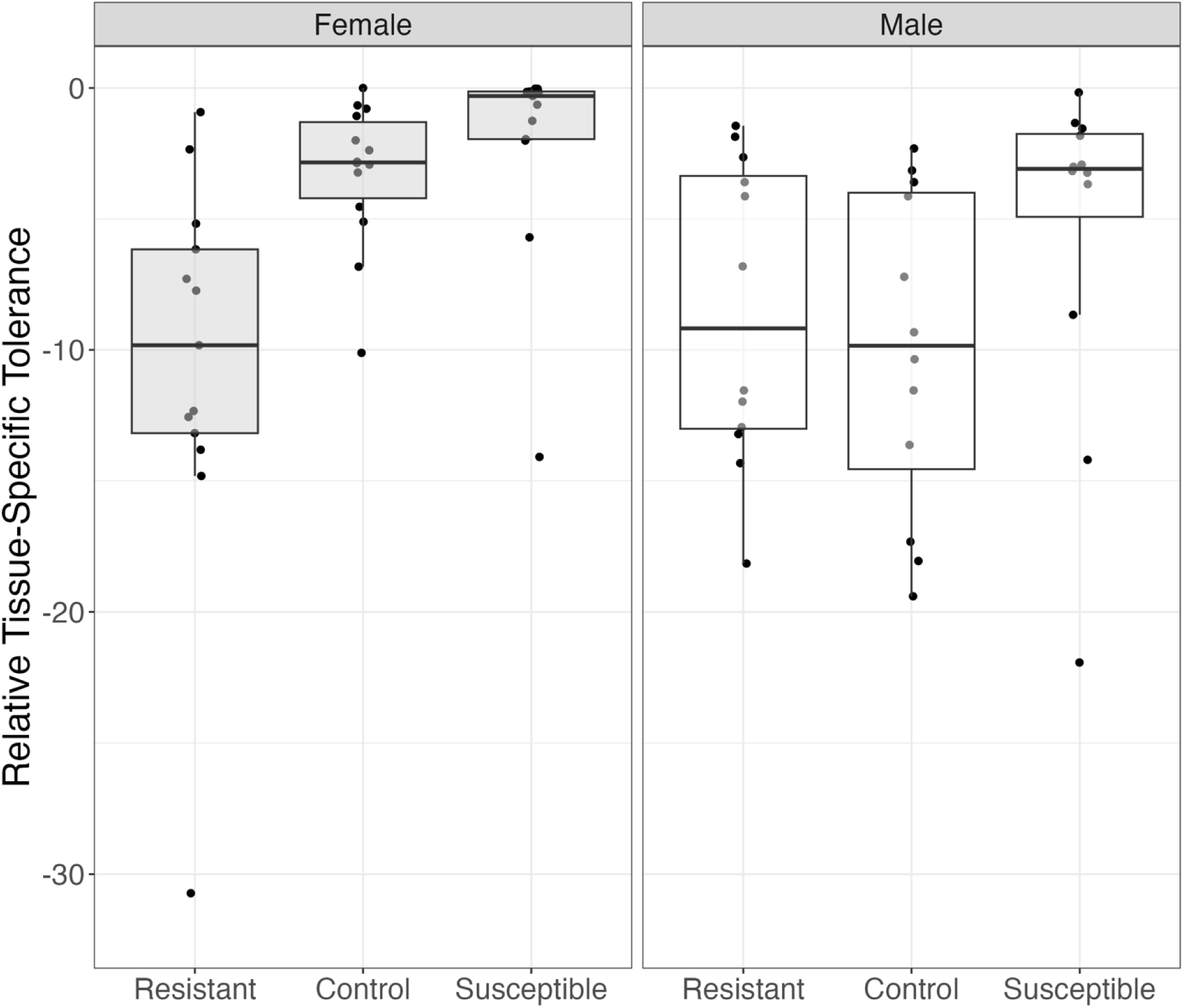
Artificial selection lines also predicted relative tissue-specific tolerance in Trinidadian Guppies. Tolerance measured as per-parasite mass change across infection minus max tolerance. Female guppies (n=44) in resistant lines significantly differed from control and susceptible lines. Male guppies (n=43) did not show significant differences. Box plots illustrate the same metrics as in Figure 1.

The polynomial regression model revealed a tradeoff between tolerance and resistance at the individual level: higher tolerance was correlated with lower resistance (Figure S1; F= 55.49, p<7e-15). No other fixed effects included explained significant variation in tolerance.

#### 3.1.3 Selection lines did not differ in the extent of inbreeding

We saw a significant reduction in allelic richness among all experimental infection groups relative to the wild-caught founders, likely reflecting a post-capture reduction in population size (Table S1). However, we found no significant difference in AR (p= 0.589), Hs (p=0.73), FIS (p=0.42), or g2 (Table S2) among the three lines.

### 3.2 The host-associated microbiome is associated with guppy resistance to Gyrodactylus turnbulli infection, independently and to a similar degree as host genetics

DMM modeling found that the lowest Laplace approximation model fit was three bacterial clusters (*Figure S2*). All samples had a posterior probability of 98% or higher. Each bacterial community type contained 10.0–66.8% of all samples (n=5-33), and no samples were undefined. The three bacterial community types (A, B, and C) in our fish skin samples were distinct (PERMANOVA: F = 7.37, p = 0.0003, Figure 3a & Figures S3-4). Males and females were balanced among the bacterial community types A and B, and bacterial community types clustered evenly across the artificial selection lines (Figure 3a; Fisher’s exact test, p=0.8). Additionally, we found no interaction between line and bacterial community types in our model (χ2(4)=3.07, p=0.546). All skin swab samples were assigned to a community type with the greatest posterior probability. For top contributing bacterial genera, see *Figure S3*.

**Figure 3.**
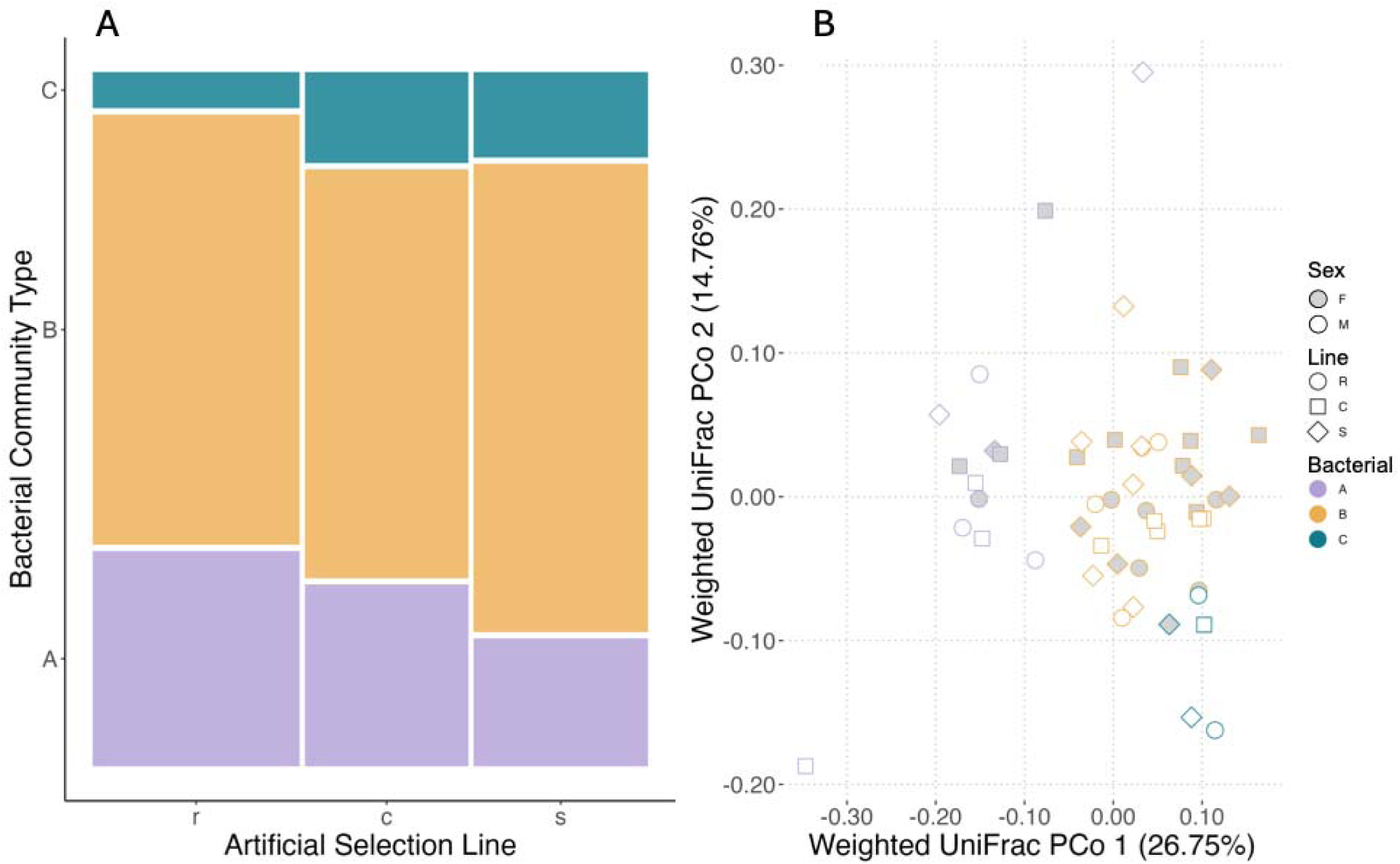
The distribution of skin bacterial community types across guppy selection lines showed no significant association. A. mosaic plot presents bacterial community types (A, B, C) across control (c), resistant (r), and susceptible (s) lines, with no significant distribution differences (Fisher’s exact test, p=0.8). B. PCoA analysis (n=52) indicated bacterial community type was significant in variation (PERMANOVA: F=7.85, P=0.001), but no significant clustering was detected for selection lines (PERMANOVA: F=0.81, P=0.95).

The selection lines did not significantly differ in any skin microbial diversity metrics tested. The microbiomes associated with females had significantly higher alpha diversity than those associated with males (Kruskal–Wallis pairwise test, Shannon’s index: H=4.06, q=0.04; observed ASVs: H=4.88, q=0.03). Consistent with Kramp et al. (2022), the microbiomes associated with the two sexes also had significantly different beta diversity (unweighted unifrac PERMANOVA; F=1.56, p=0.006).

We found that the bacterial community type A of the skin microbiome on Trinidadian guppies had significantly reduced infection severity compared to bacterial community type B & C (χ^2^(2) = 19.83, p=4.9e-5, Figure 3b). A post-hoc test showed that a model including both selection line and bacterial community type is significantly better at explaining variation in infection severity than models with either variable alone (Figure 4; χ^2^(14)=13.40, p=2e-16).

**Figure 4.**
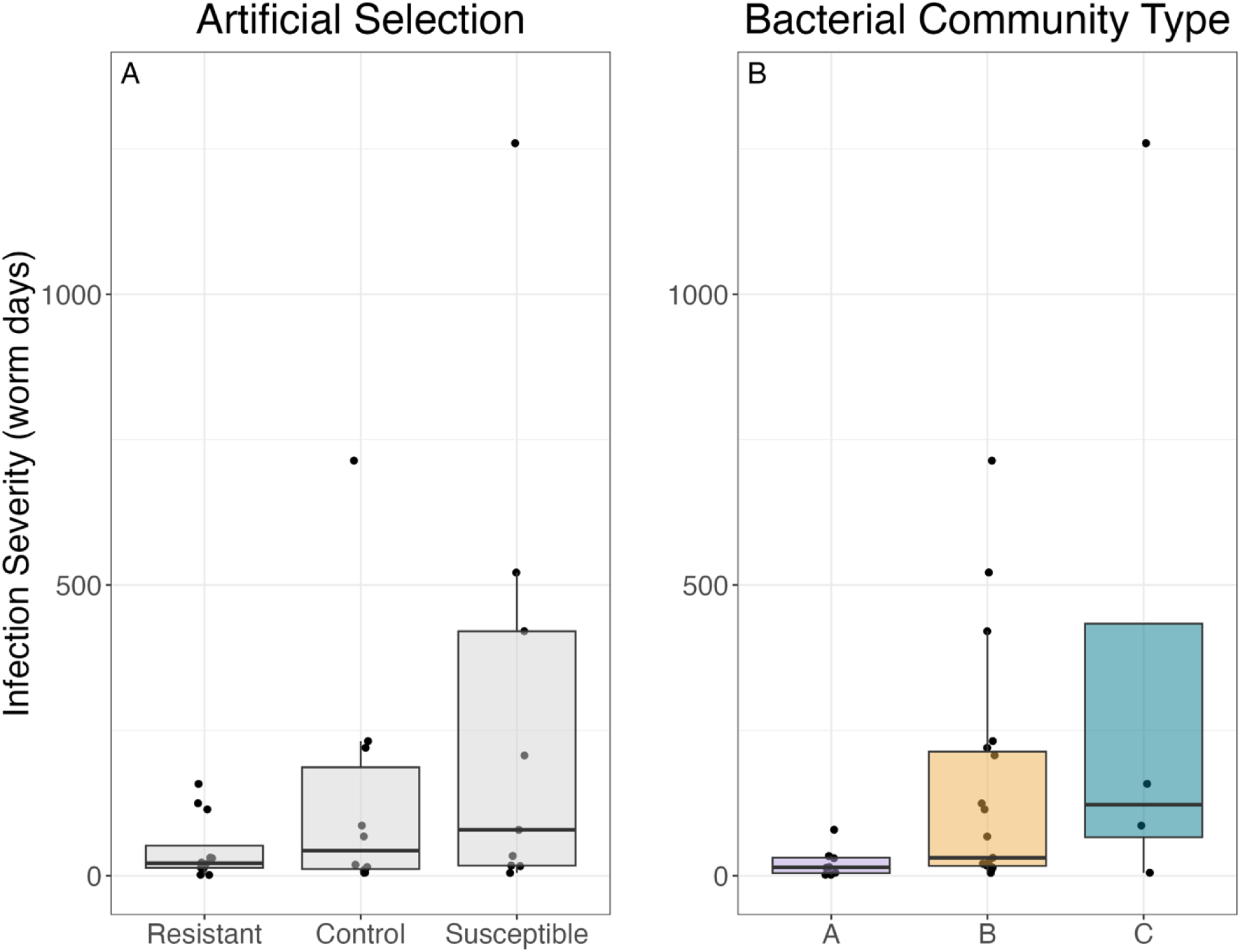
Selection lines and bacterial community types predicted G. turnbulli infection severity. A. Lines significantly predicted severity in swabbed fish (n=31; ANOVA: χ²=8.73, P=0.013). B. Community type A had lower infection severity than types B and C (A-B, z=3.23, P=0.0034; A-C, z=4.92, P<0.001). No interactions between community type and selection line were detected (ANOVA: χ²=7.4, P=0.116). A combined model of line and bacterial types explained infection severity better than either factor alone (χ²=13.40, P<0.001).

## 4. Discussion

Our selection lines demonstrated striking heritability of parasite infection resistance and tolerance. Despite no significant differences in microbiome between the three lines, we found that bacterial community type, found by DMM machine-learning, predicted infection severity. Host genetics and the host-associated microbiome thus appear to independently predict infection severity, with similar effect sizes. We also observed notable differences between males and females in skin microbiome, and in response to selection. We discuss each of these results in turn.

We found that both guppy resistance and susceptibility to *G. turnbulli* responded readily to a single generation of truncation selection. We found no notable differences between the three selection lines in our genetic metrics, indicating that the differences we found were not due to differences in inbreeding among them. Our narrow-sense heritability estimates for resistance and tolerance were high, particularly for our susceptible lines. However, our values are comparable to heritability estimates in previous studies (Weiler et al., 2025) and to the heritability of male guppy ornaments (Morris et al., 2020), which are correlated with *G. turnbulli* resistance (Stephenson et al., 2020).

Parasite resistance and susceptibility likely responded to selection through different mechanisms. The response to selection for susceptibility could have involved either an increase in tolerance, a loss of resistance, or some combination. Indeed, our experimental design may have favored tolerance in the susceptible line, as only individuals surviving severe infections would have contributed to future generations. Tolerance and resistance likely have a different genetic bases, and hosts appear to invest in one strategy at the expense of the other (Råberg et al., 2007). This observation is supported by the fact that our resistant line fish had lower tolerance than control and susceptible line fish – at least among females – and that the two defense strategies were negatively correlated at the level of individual fish in our experiment. However, these tradeoffs do not always occur, and both tolerance and resistance traits can coexist within a single individual host and a population (Hardy et al., 2024; Pagán & García-Arenal, 2018).

Our observed high heritability of susceptibility (and potentially tolerance) has implications for our understanding of how guppy defenses might evolve in natural conditions (Walsman et al., 2022, 2024). High heritability suggests natural selection can act rapidly on host defense traits in populations under high parasite pressure, and indeed host defense appears to have the capacity to evolve rapidly in this system (Dargent et al., 2016; Phillips et al., 2018). In some populations, infection prevalence is very high (>90%; (Clark et al., 2023; Stephenson, van Oosterhout, Mohammed, et al., 2015)), potentially selecting for increased infection tolerance and decreased resistance, a phenomenon known as the ‘resistance is futile’ effect (Walsman et al., 2023). However, we (and all previous studies of host defense traits in this system) deliberately excluded any potential role of parental effects of infection, when recent findings suggest that transgenerational immune priming may be important in this system, which would reduce the strength of selection on these traits, particularly when prevalence is high (Weiler et al., 2025).

We found that bacterial community type predicted infection severity (but interestingly not tolerance) independently, and to a similar degree, as host genetics. Despite the fact that host characteristics such as the immune system, metabolism, and growth rate have been shown to interact with the microbiome (Speer, 2022; Zheng et al., 2020), we found no difference between our lines in skin microbiome composition, alpha diversity, or bacterial community types. Potentially this is because fish skin microbiomes are highly variable and may not reflect the host’s genetics but instead drift stochastically (Jones et al., 2022; Wang et al., 2023). However, recently it has come to light that different organ may provide unique insights of host health, i.e., eye microbiome predicted disease outcome but gut microbiome did not in Island foxes (DeCandia et al., 2025). Therefore, it is possible that microbiome from other tissue could have been modified due to our selection, though, we did not test for this. By contrast, artificial selection for body size in gilthead sea bream (*Sparus aurata*) significantly altered the host-associated microbiome, and the microbiome in the fast-growing line had greater parasite colonization resistance (Piazzon et al., 2020), and size selection experiments indirectly affect guppy resistance to *G. turnbulli* (Bartuseviciute et al., 2022) potentially mediated through the microbiome. These results suggest that selection on host growth does influence the associated microbiome composition, potentially through effects on host metabolism, with downstream effects on parasite resistance (Bartuseviciute et al., 2022; Piazzon et al., 2020), whereas selection on parasite resistance appears not to have direct effects on skin-associated microbial communities (this study).

Our findings thus contribute to the growing literature linking host-associated microbiomes and infection severity, pointing to important interactions between the skin microbiome and parasites (Scheifler et al., 2022; Speer, 2022; Trevelline et al., 2020). More broadly, the microbiome’s role in determining resistance has been seen in several taxa, where pre-infection microbiome composition, alpha diversity, or microbiome function is a powerful tool to predict individuals that are most susceptible to parasites within a population (Bates et al., 2022; Lee et al., 2019). Intriguingly, we found five of the eight ASVs reported to be indicator taxa of epizootic dynamics in a recent study of skin microbial metabolism function in amphibian chytridiomycosis (Bates et al., 2022): *Burkholderiales*, *Pseudomonadales*, *Sphingomonadales*, *Rhizobiales*, *Flavobacteriales* (see Figure S2 for all contributing species).

Sex differences are pervasive throughout our results. Sexual dimorphism in immunocompetence has been reported in many taxa, including Trinidadian guppies. Our results reveal additional evidence of sex-specificity in defenses against the specialist parasite *G. turnbulli*: resistance responded more strongly to selection among female than male guppies, as has been found previously (Dargent et al., 2016). Potentially, this result reflects the fact that males in nature and in our experiment were subject to sexual selection in addition to direct selection for parasite resistance. Females in many populations prefer males with larger orange ornaments (Endler et al., 2001; Houde, 1988, 1997), with implications for disease dynamics (Rovenolt et al., 2024). Larger orange ornaments indicate male resistance to this parasite (Stephenson et al., 2020). Males tend to develop lower infection loads in general in the lab (Johnson et al., 2011), are less likely, and less heavily infected in surveys of natural populations (Clark et al., 2023; Gotanda et al., 2013; Stephenson, van Oosterhout, Mohammed, et al., 2015). Thus, if female preference has already eroded genetic variation in parasite resistance among males, and females across our lines selected the most resistant males as mates, this may explain the overall high and invariable resistance we observe in males. By contrast, and again in line with previous results from field surveys (Stephenson, van Oosterhout, & Cable, 2015), females – particularly large ones – appear to be more tolerant of infection than males. Notably, we found evidence for a tradeoff between resistance and tolerance at the line level among females, but not among males, perhaps indicating different trade-offs for each sex, though there was no sex difference in the tradeoff at the level of individual fish.

The strong sex-specificity in host defense traits we observe here highlights the need to investigate aspects of guppy response to *Gyrodactylus turnbulli* beyond the MHC. Specifically, if the primary mechanism of resistance were driven by MHC parasite resistance, we would not see such pronounced sex differences in response to selection: MHC resistance is a mixture of both mother and father MHC genes. While our data indicate that the microbiome is not driving this sex difference in response to selection, the microbiome clearly contributes to parasite resistance in this system and appears to differ between the sexes: they differed in both alpha and beta diversity metrics of bacterial communities, similar to the skin microbiome in other species (Rodríguez-Barreto et al., 2023). The observed sex-specific differences highlight potential co-evolutionary interactions between host sex and microbiome in relation to parasite resistance. These possibilities will hopefully inspire exciting and novel exploration of the evolutionary dynamics of resistance and tolerance between the sexes.

## Author Contributions

Rachael Kramp and Jessica Stephenson conceived the ideas and designed methodology; Jessica Stephenson and Nadine Tardent collected the data; Rachael Kramp, Jessica Stephenson and Mary J. Janecka analysed the data; Rachael Kramp, Jessica Stephenson, Kevin Kohl and Jukka Jokela led the writing and editing of the manuscript. All authors contributed critically to the drafts and gave final approval for publication.

## Animal use ethics statement

The permit to collect and export guppies was granted by the Director of Fisheries in the Ministry of Agriculture, Land and Fisheries Division, Aquaculture Unit of the Republic of Trinidad and Tobago. Import to Cardiff University was under Cefas APB authorisation number CW054-D—187A. Fish husbandry and experimental infections of the F1 generation at Cardiff were under UK Home Office licence PPL 302876, with approval by the Cardiff University Animal Ethics Committee. Guppies were shipped to Zürich, and their husbandry and experimental infections of the F3-7 generations were approved by the Veterinäramt of Kanton Zürich (ZH177/15).

## Data availability statement

The raw experimental data, description of the data and analysis code are available from the Dryad Digital Repository: http://datadryad.org/share/UUQY9h2hXgGiWvvPM_SC6D2ii3ssYFAq71k3w4dFKlI

