## Supplemental Figures for "Host genetics and the Skin Microbiome Independently Predict Parasite Resistance"

### Artificial Selection and the Skin Microbiome Independently Predict Parasite Resistance in Poecilia reticulata

anonymous review

**4 Part II: Looking deeper:The host-associated microbiome clustering in artifical selection lines or independently coorelating with infections integral** **28**

5 PART III: Investigation of explanatory power between artifical selection and microbiome. 37

### 1 Introduction

In this study, we wanted to understand how certain factors can affect the outcome of parasitic infections. These factors include artificial selection, the sex of the fish, and the community of microorganisms living on the fish's skin.

To set up the artificial selection process, we infected 200 *Poecilia reticulata* fish (half male and half female) with a parasite called *Gyrodactylus turnbulli*. We closely monitored the infection, then divided the fish into two groups: one group with the top 30% of fish having the highest average worm burden (susceptible line) and the other group with the bottom 30% of fish having the lowest average worm burden (resistance line). We also had control groups that were made up of uninfected fish randomly selected from the same population.

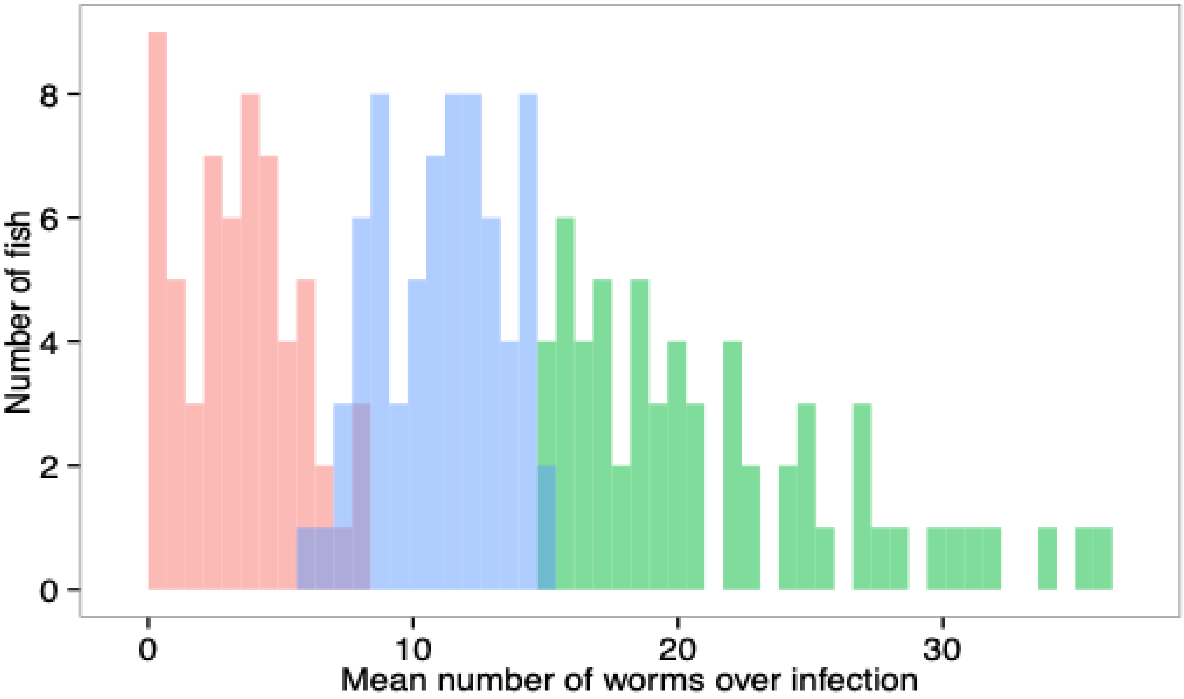

Figure 1: Histogram of Individuals chosen for Artificial selection "parental population" of *poecilia reticulata* for resistance and susceptibility to *Gyrodactylus turnbulli*. A Histogram showing individual fish frequency distribution of mean number of *G. turnbulli* parasites or "worms" during infection of parental lines. Fish (n=200). Pink color represents the 30% least intense infection loads and fish were used to establish "Resistant line". Green color represents the 30% most intense infection loads and fish were used to establish "susceptible line". Blue color is excluded fish.

After this initial selection process, we allowed the fish from each group to breed within their respective lines for four to seven generations without exposure to the parasite. Before infecting the offspring from the artificial selection, we collected swabs from a subset of the adult fish to measure the communities of microorganisms living on their skin.

In our analysis, the most important data can be found in the file 'SelectionData.csv,' which contains various variables used in our main statistical models:

- **fishID:** Identity of the focal fish
- **line:** The 30% most intense infection founded the susceptible line and the 30% least intense infections founded the resistance lines. The controls lines were founded with the same number of parents of randomly selected uninfected fish from the same population.
- **JulienDays:** Experimental dates when the fish were swabbed and infected
- **sex:** Sex of the fish
- **length, weight:** Length in mm and weight (mass) in mg of the fish
- **respreL:** The residuals of length of the focal fish on sex. Females tend to be larger than males: using the residuals of this relationship allows us to test for both size and sex differences in behavior.
- **PREsmi** The residuals of the scaled mass index of fish
- **gen** The generation the fish was assigned to
- **dose** The number of parasites that jumped onto the experimental fish during infection or "dose"
- **rSMIpre** Body condition - (rescaled so that males and females could be compared) scaled mass index
- **AUC** The Area Under the Curve for worm days / time "infection integral"
- **maxworm** The maximum number of parasites recorded on that fish during its infection
- **meanworm** average number of worms or parasite individual counted on the guppy from day 1-9 of infection
- **pmass** Percentage mass change over the course of infection
- **rscmass**rescaled all %body mass changes relative to the highest %body mass
- **ppmass** using "rscmass" or we divded by maxworm to get our "relative tissue-tolerance"

This data was subsetting to run analysis appropriately, all dataframes are explained below.

The following sub-setting of Dataframe's explained:

**Parental Data frame**: Only founders of the R,S lines to look at heriability metrics within the resistant and suscepitble lines

**IOm Data frame**: Only offspring that were infected with *G.turnbulli*, and will be used in GLMM model.

**female and male Data frames**: Subset F4-7 offspring fish that were infected with *G. turnbulli* parasite sub-setting to contain only female fish or male fish for post-hoc analysis

### 2 Visualization of offspring data

We used pairwise plots to check distribution of single variables, multicollinearity, and correlations between our variables of interest.

```r
options(width = 70, digits = 22)

## running pairs plot for all variables

pairs(~AUC + sex + rPREsmi + respreL + JulianDate + line + pmass + rscmass +
    ppmass + maxworm + dose + gen, data = IOm, lower.panel = panel.smooth,
    upper.panel = panel.cor, na.action = na.omit)
```

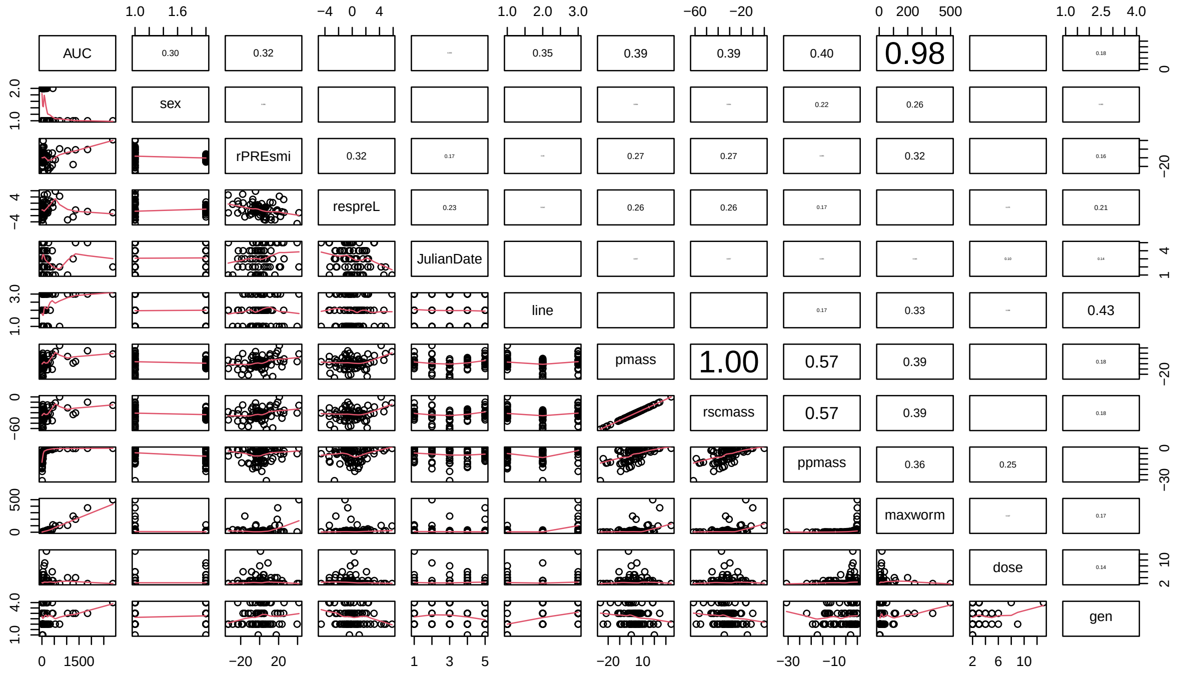

### 3 Part I: Artificial selection for parasite resistance in Trinidadian Guppies

#### 3.1 Testing for differences between the A and B subpopulations within each line

Here we evaluate whether there are differences between the A and B subpopulations within each line. Each line was divided in F2 (i.e., first generation post selection) to control for tank effects (note that after F3 each line was held in several different tanks) and other stochastic factors that might affect fish during generations of breeding. The analysis below shows no difference between the A and B subpopulations within each line in either AUC or ppmass, so we combined the subpopulations in the next steps of the analysis.

```r
# create variable combining line and rep for visualisations
IOm$lr <- paste(IOm$line, IOm$rep, sep = ".")
class(IOm$lr)
```

```
[1] "character"
```

```
IOm <- as.data.frame(IOm)

# do the reps differ in AUC?
TMBModel <- glmmTMB(AUC ~ line * rep, data = IOm, family = Gamma(link = "log"))
Anova(TMBModel)  #used for using P-values
```

```
Analysis of Deviance Table (Type II Wald chisquare tests)

Response: AUC
                         Chisq Df Pr(>Chisq)
line     42.98319000000000004525  2 4.6379e-10 ***
rep       1.07787000000000001058  1    0.29917
line:rep  2.22251999999999998290  2    0.32914
---
Signif. codes:
  0 '***' 0.00100000000000000020817 '**' 0.01000000000000000020817
  '*' 0.05000000000000000277556 '.' 0.10000000000000000055511 ' ' 1
```

```
# visualization of results and verifying model with residuals
# Diagnostic plots
sim_residuals_auc <- simulateResiduals(TMBModel, 1000)
plot(sim_residuals_auc)
```

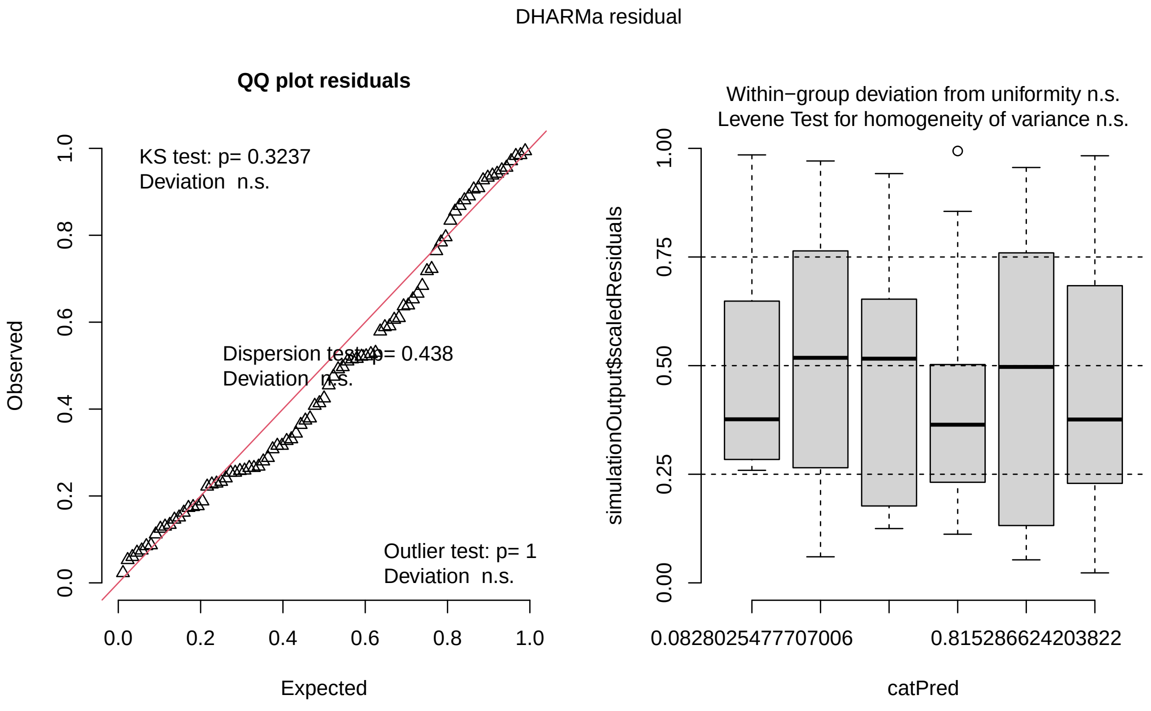

```
testDispersion(sim_residuals_auc)
```

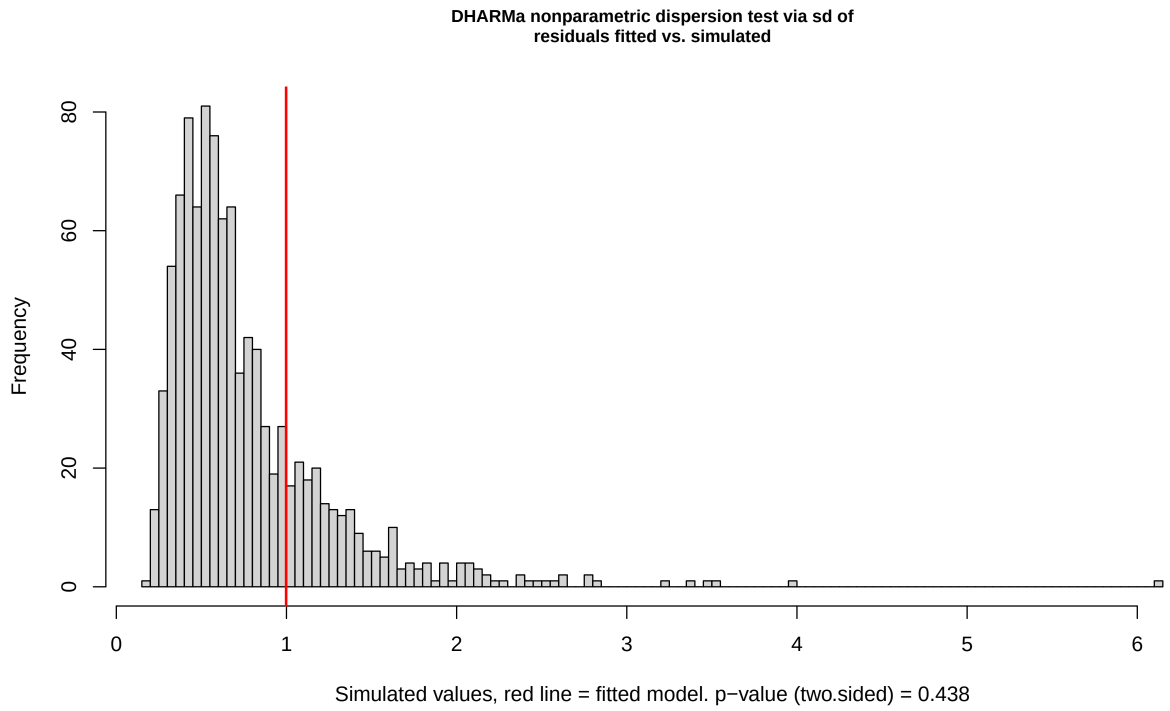

```
    DHARMa nonparametric dispersion test via sd of residuals
    fitted vs. simulated

data:  simulationOutput
dispersion = 1.2939155249104115075, p-value =
0.43800000000000000004
alternative hypothesis: two.sided
```

```
# do the reps differ in ppmass?
TMBModel1 <- glmmTMB(ppmass ~ line * rep, data = IOm)

Anova(TMBModel1)  #used for using P-values
```

```
Analysis of Deviance Table (Type II Wald chisquare tests)

Response: ppmass
                          Chisq Df Pr(>Chisq)
line     12.7146500000000006736  2   0.001734 **
rep       2.8148800000000000487  1   0.093394 .
line:rep  1.8145599999999999508  2   0.403621
---
Signif. codes:
  0 '***' 0.00100000000000000020817 '**' 0.01000000000000000020817
  '*' 0.05000000000000000277556 '.' 0.10000000000000000555111 ' ' 1
```

```
# visualization of results and verifying model with residuals
# Diagnostic plots
```

```r
sim_residuals_auc <- simulateResiduals(TMBModel1, 1000)
plot(sim_residuals_auc)
```

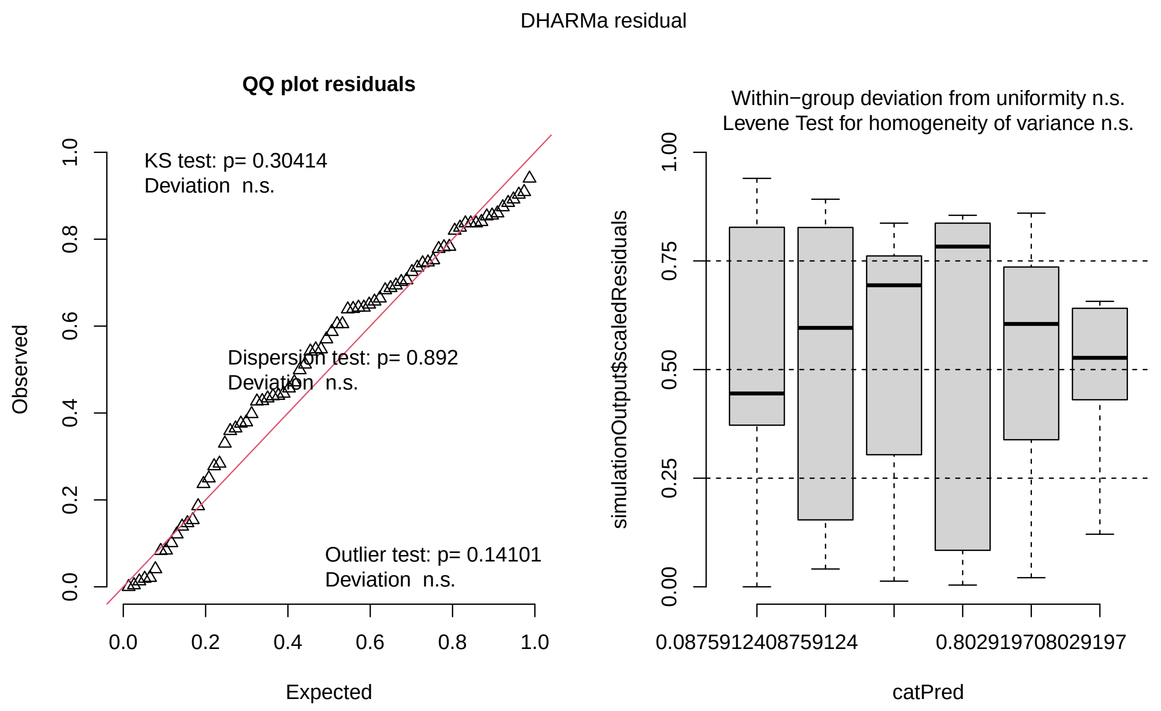

```r
testDispersion(sim_residuals_auc)
```

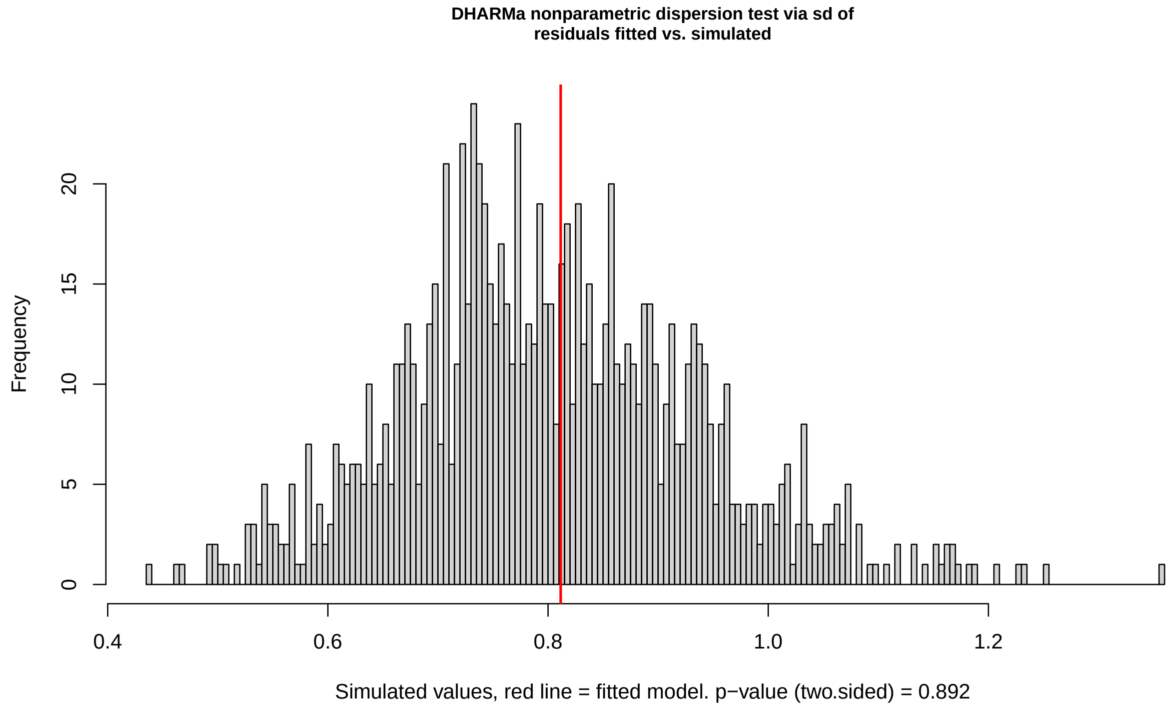

```
    DHARMa nonparametric dispersion test via sd of residuals
    fitted vs. simulated

data:  simulationOutput
dispersion = 1.0102157455955758092, p-value =
0.89200000000000000151
alternative hypothesis: two.sided
```

##### 3.2 Generalized linear mixed model (GLMM model): Variables explaining infection infectional intrgral after imposed Artifical Selection (generations 4-7)

Generations 4-7 of Artificial Selection were infection with *G. turnbulli* and infection was quantified for 14 days. Did the Artificial selection work, do we see infection integral differences between susceptible, resistant, and control fish?

**Response variable:** the area under the curve for worm days/ time "infection integral".

**Fixed Effects:**

- Line (F4-7 Resistant, susceptible, control fish)
- rPREsmi (residuals of scaled mass index "body condition" of fish)
- respreL (residual of fish lengths)
- Sex of fish
- Julian Date (Date fish were infected)

```r
options(width = 70, digits = 4)

# When controlling for SMI, length, generation time, and total
# starting parasite numbers is there infection differences in R,S,C
# fish?

TMBModel <- glmmTMB(AUC ~ rPREsmi + respreL + gen + dose + line * sex +
    JulianDate, data = IOm, family = Gamma(link = "log"))

Anova(TMBModel)  #used for using P-values
```

```
Analysis of Deviance Table (Type II Wald chisquare tests)

Response: AUC
           Chisq Df Pr(>Chisq)
rPREsmi     0.35  1     0.5560
respreL     3.62  1     0.0569 .
gen         1.14  3     0.7683
dose        5.11  1     0.0238 *
line       37.74  2    6.4e-09 ***
sex        24.29  1    8.3e-07 ***
JulianDate  9.48  4     0.0501 .
line:sex   13.51  2     0.0012 **
---
Signif. codes:  0 '***' 0.001 '**' 0.01 '*' 0.05 '.' 0.1 ' ' 1
```

```r
summary(glht(TMBModel, linfct = mcp(line = "Tukey")))
```

```
Warning in mcp2matrix(model, linfct = linfct): covariate interactions
found -- default contrast might be inappropriate

     Simultaneous Tests for General Linear Hypotheses

Multiple Comparisons of Means: Tukey Contrasts

Fit: glmmTMB(formula = AUC ~ rPREsmi + respreL + gen + dose + line *
    sex + JulianDate, data = IOm, family = Gamma(link = "log"),
    ziformula = ~0, dispformula = ~1)

Linear Hypotheses:
           Estimate Std. Error z value Pr(>|z|)
c - r == 0    0.739      0.402    1.84     0.16
s - r == 0    3.105      0.524    5.92   <1e-04 ***
s - c == 0    2.365      0.471    5.02   <1e-04 ***
---
Signif. codes:  0 '***' 0.001 '**' 0.01 '*' 0.05 '.' 0.1 ' ' 1
(Adjusted p values reported -- single-step method)
```

```r
# visualization of results and verifying model with residuals
# Diagnostic plots
sim_residuals_auc <- simulateResiduals(TMBModel, 1000)
plot(sim_residuals_auc)
```

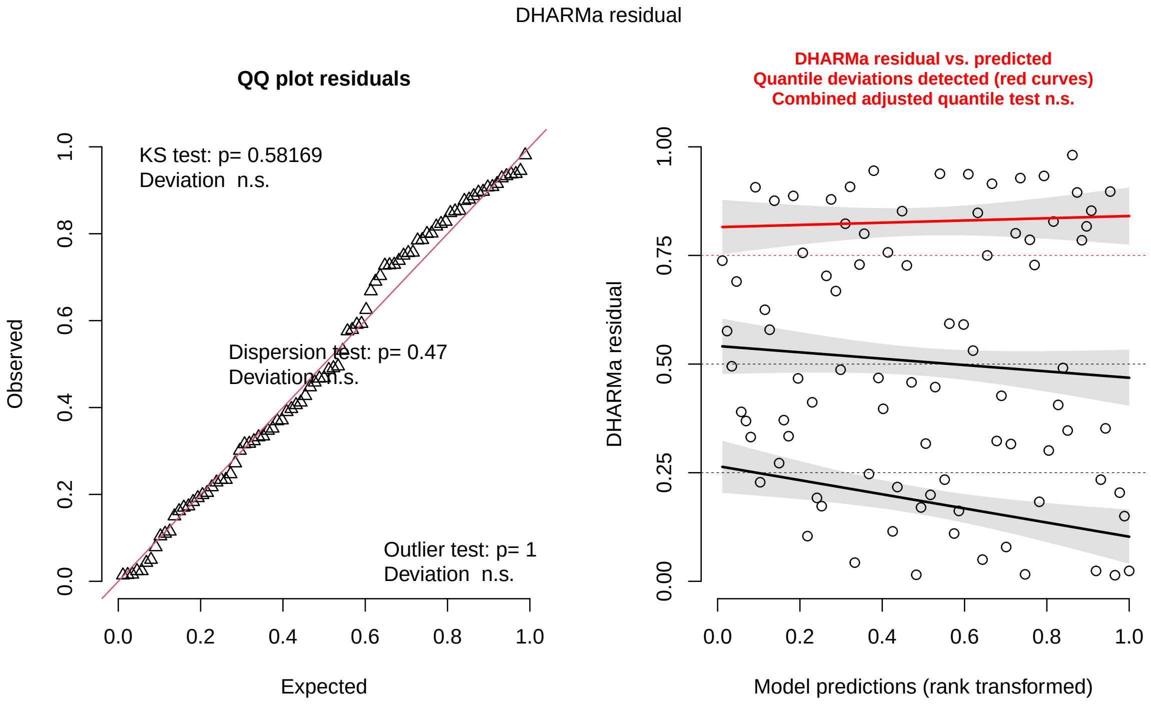

```
testDispersion(sim_residuals_auc)
```

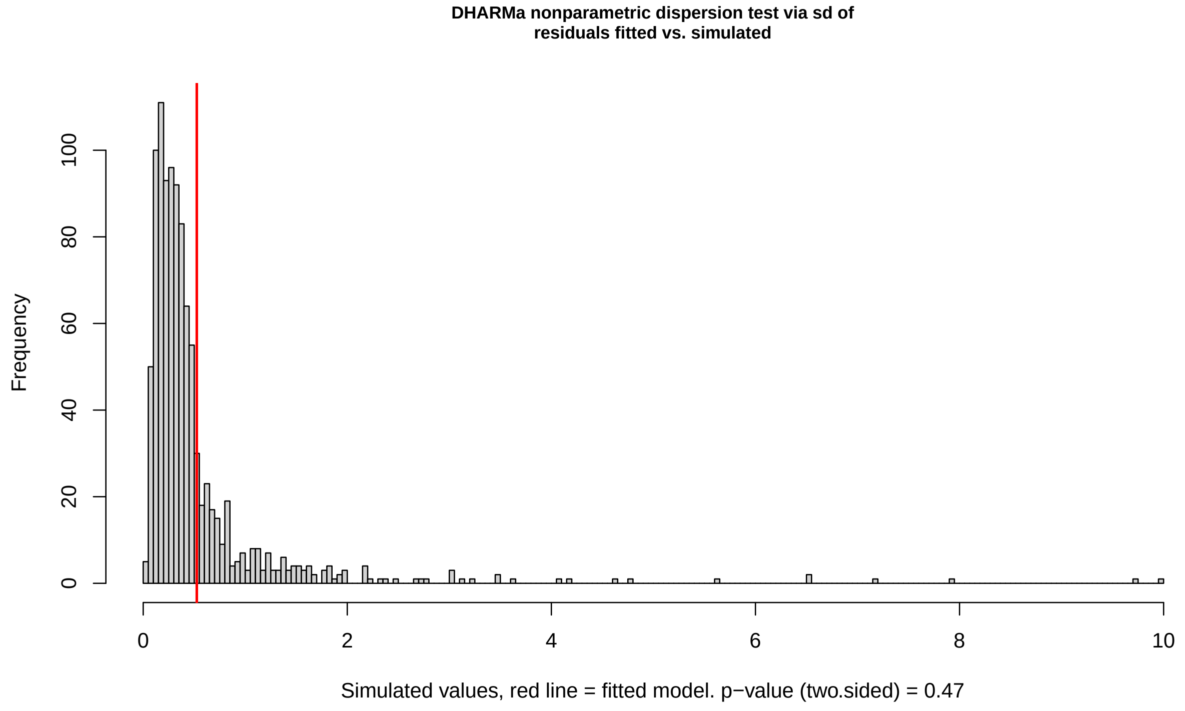

```
    DHARMa nonparametric dispersion test via sd of residuals
    fitted vs. simulated

data:  simulationOutput
dispersion = 1, p-value = 0.5
alternative hypothesis: two.sided
```

##### 3.3 Sex Post-hoc analysis GLMM: The correlation between the artificial selection line and infection integral is being assessed for significant differences from 0 in either sex.

Due to the interaction between our 'line' and 'sex' of fish we preformed a post-hoc analysis on male and female fish to see if only female or male fish responded to artificial selection.

```
options(width = 70, digits = 4)
# Do females have the same results as seen above?

FeTMB <- glmmTMB(AUC ~ line + rPREsmi + respreL + gen + dose + JulianDate,
    data = female, family = Gamma(link = "log"))
## stats for model
Anova(FeTMB)
```

```
Analysis of Deviance Table (Type II Wald chisquare tests)
```

```
Response: AUC
           Chisq Df Pr(>Chisq)
line       24.33  2    5.2e-06 ***
rPREsmi     1.09  1      0.295
respreL     2.90  1      0.089 .
gen         1.28  3      0.734
dose        0.77  1      0.381
JulianDate  4.99  4      0.288
---
Signif. codes:  0 '***' 0.001 '**' 0.01 '*' 0.05 '.' 0.1 ' ' 1
```

```r
summary(glht(FeTMB, linfct = mcp(line = "Tukey")))
```

```
     Simultaneous Tests for General Linear Hypotheses

Multiple Comparisons of Means: Tukey Contrasts

Fit: glmmTMB(formula = AUC ~ line + rPREsmi + respreL + gen + dose +
    JulianDate, data = female, family = Gamma(link = "log"),
    ziformula = ~0, dispformula = ~1)

Linear Hypotheses:
           Estimate Std. Error z value Pr(>|z|)
r - c == 0   -1.129      0.412   -2.74   0.0162 *
s - c == 0    1.919      0.550    3.49   0.0011 **
s - r == 0    3.049      0.620    4.92   <0.001 ***
---
Signif. codes:  0 '***' 0.001 '**' 0.01 '*' 0.05 '.' 0.1 ' ' 1
(Adjusted p values reported -- single-step method)
```

```r
# visualization of results and verifying model with residuals
# Diagnostic plots
sim_residuals_auc <- simulateResiduals(FeTMB, 1000)
plot(sim_residuals_auc)
```

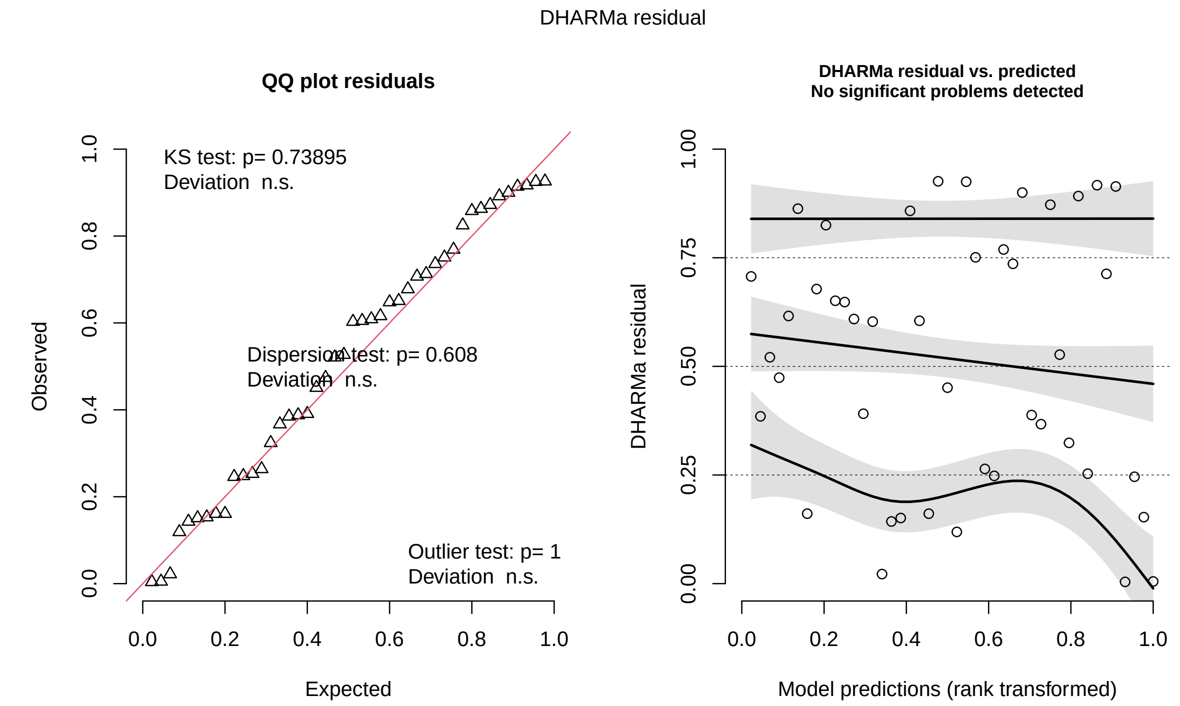

```r
testDispersion(sim_residuals_auc)
```

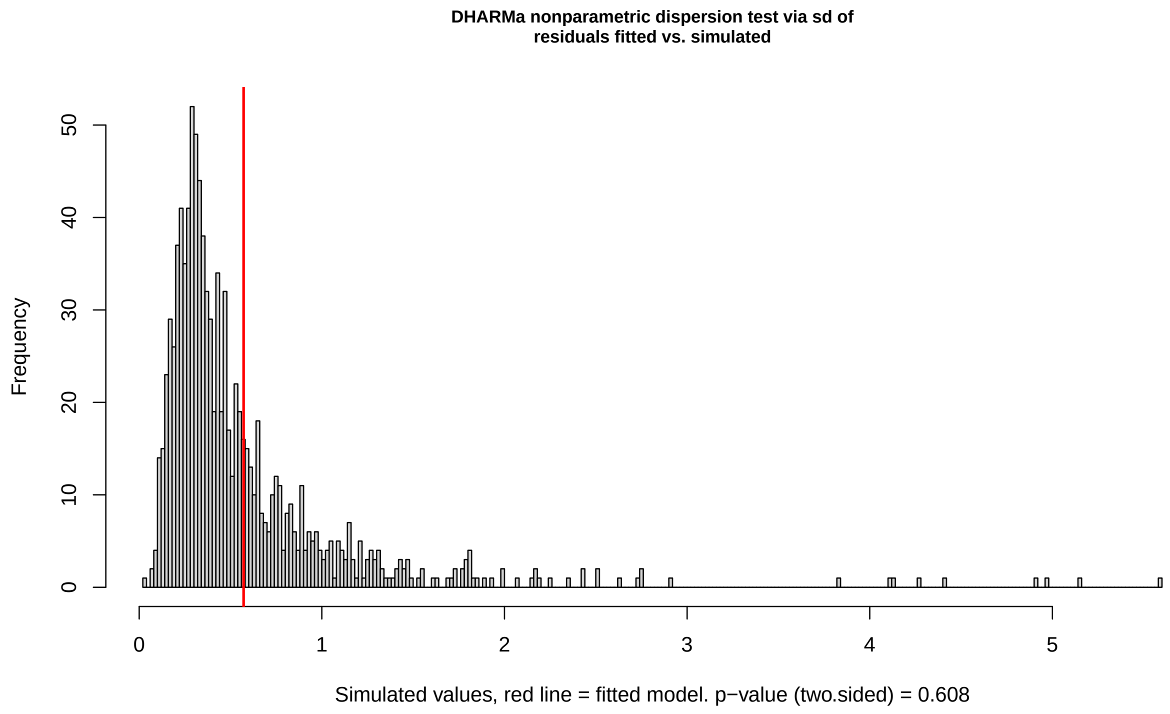

```
DHARMa nonparametric dispersion test via sd of residuals
```

```
    fitted vs. simulated

data:  simulationOutput
dispersion = 1, p-value = 0.6
alternative hypothesis: two.sided
```

```
# Do males have the same results as seen above?

maTMB <- glmmTMB(AUC ~ line + rPREsmi + respreL + gen + dose + JulianDate,
    data = male, family = Gamma(link = "log"))

## stats for model
Anova(maTMB)
```

```
Analysis of Deviance Table (Type II Wald chisquare tests)

Response: AUC
           Chisq Df Pr(>Chisq)
line       17.57  2    0.00015 ***
rPREsmi     0.01  1    0.92680
respreL     0.01  1    0.90268
gen         0.55  3    0.90687
dose        7.88  7    0.34337
JulianDate  2.64  4    0.62050
---
Signif. codes:  0 '***' 0.001 '**' 0.01 '*' 0.05 '.' 0.1 ' ' 1
```

```
summary(glht(maTMB, linfct = mcp(line = "Tukey")))
```

```
     Simultaneous Tests for General Linear Hypotheses

Multiple Comparisons of Means: Tukey Contrasts

Fit: glmmTMB(formula = AUC ~ line + rPREsmi + respreL + gen + dose +
    JulianDate, data = male, family = Gamma(link = "log"), ziformula = ~0,
    dispformula = ~1)

Linear Hypotheses:
           Estimate Std. Error z value Pr(>|z|)
r - c == 0    0.896      0.483    1.86    0.151
s - c == 0    2.288      0.546    4.19   <0.001 ***
s - r == 0    1.392      0.529    2.63    0.023 *
---
Signif. codes:  0 '***' 0.001 '**' 0.01 '*' 0.05 '.' 0.1 ' ' 1
(Adjusted p values reported -- single-step method)
```

```
# visualization of results and verifying model with residuals
# Diagnostic plots
sim_residuals_auc <- simulateResiduals(maTMB, 1000)
plot(sim_residuals_auc)
```

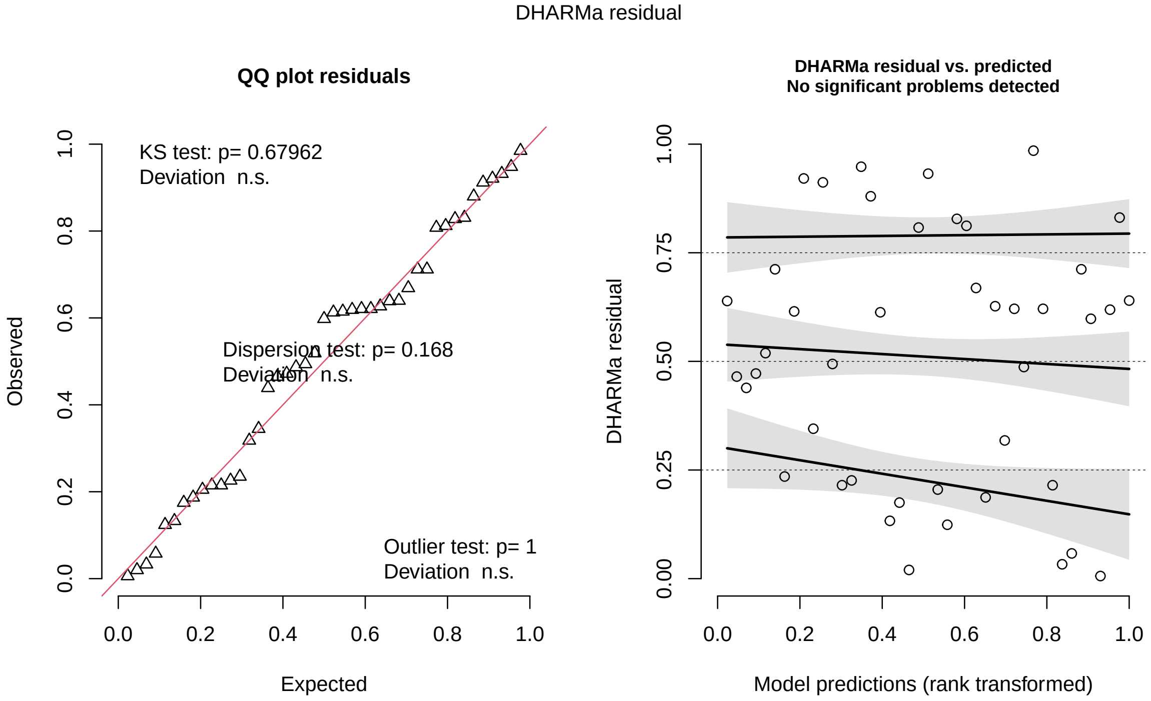

```r
testDispersion(sim_residuals_auc)
```

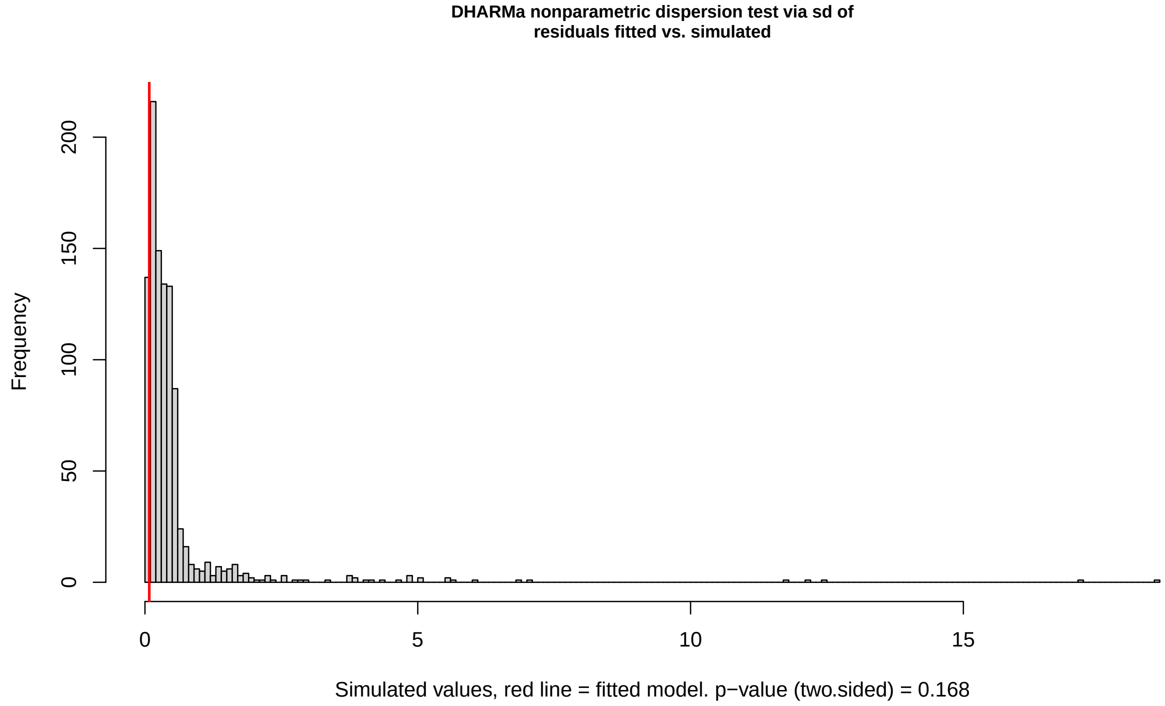

```
DHARMa nonparametric dispersion test via sd of residuals
```

```
	fitted vs. simulated

data:  simulationOutput
dispersion = 0.14, p-value = 0.2
alternative hypothesis: two.sided
```

##### 3.4 Tissue-specific tolerance: does per-parasite percentage body condition (scaled mass index) change "Tissue-specific tolerance" differ across the Artifical Selection Lines?

We quantified tissue-specific tolerance as the per-parasite precentage mass change across infection. We did this by taking relative percentage mass change over the course of infection divided by the maximum amount of worms observed on the fish during its infection. Higher values of this metric denote higher tissue-specific tolerance. I.e., even with high parasite loads you didn't loss a lot of weight.

```
ppModel <- glmmTMB(ppmass ~ line * sex + respreL + gen + dose + JulianDate,
    data = IOm, family = gaussian(link = "identity"), na.action = na.omit)

# Stats for Model
Anova(ppModel)
```

```
Analysis of Deviance Table (Type II Wald chisquare tests)

Response: ppmass
           Chisq Df Pr(>Chisq)
line       14.56  2    0.00069 ***
sex         3.17  1    0.07491 .
respreL     5.94  1    0.01479 *
gen         2.28  3    0.51609
dose        3.27  1    0.07073 .
JulianDate  5.07  4    0.28063
line:sex   10.28  2    0.00586 **
---
Signif. codes:  0 '***' 0.001 '**' 0.01 '*' 0.05 '.' 0.1 ' ' 1
```

```
AIC(ppModel)
```

```
[1] 488.7
```

```
summary(glht(ppModel, linfct = mcp(line = "Tukey")))
```

```
Warning in mcp2matrix(model, linfct = linfct): covariate interactions
found -- default contrast might be inappropriate

	 Simultaneous Tests for General Linear Hypotheses

Multiple Comparisons of Means: Tukey Contrasts
```

```
Fit: glmmTMB(formula = ppmass ~ line * sex + respreL + gen + dose +
    JulianDate, data = IOm, family = gaussian(link = "identity"),
    na.action = na.omit, ziformula = ~0, dispformula = ~1)

Linear Hypotheses:
           Estimate Std. Error z value Pr(>|z|)
c - r == 0     7.06       1.97    3.58    0.001 **
s - r == 0     8.22       2.18    3.78   <0.001 ***
s - c == 0     1.16       2.28    0.51    0.866
---
Signif. codes:  0 '***' 0.001 '**' 0.01 '*' 0.05 '.' 0.1 ' ' 1
(Adjusted p values reported -- single-step method)
```

```
# visualization of results and verifying model with residuals
# Diagnostic plots
sim_residuals_auc <- simulateResiduals(ppModel, 1000)
plot(sim_residuals_auc)
```

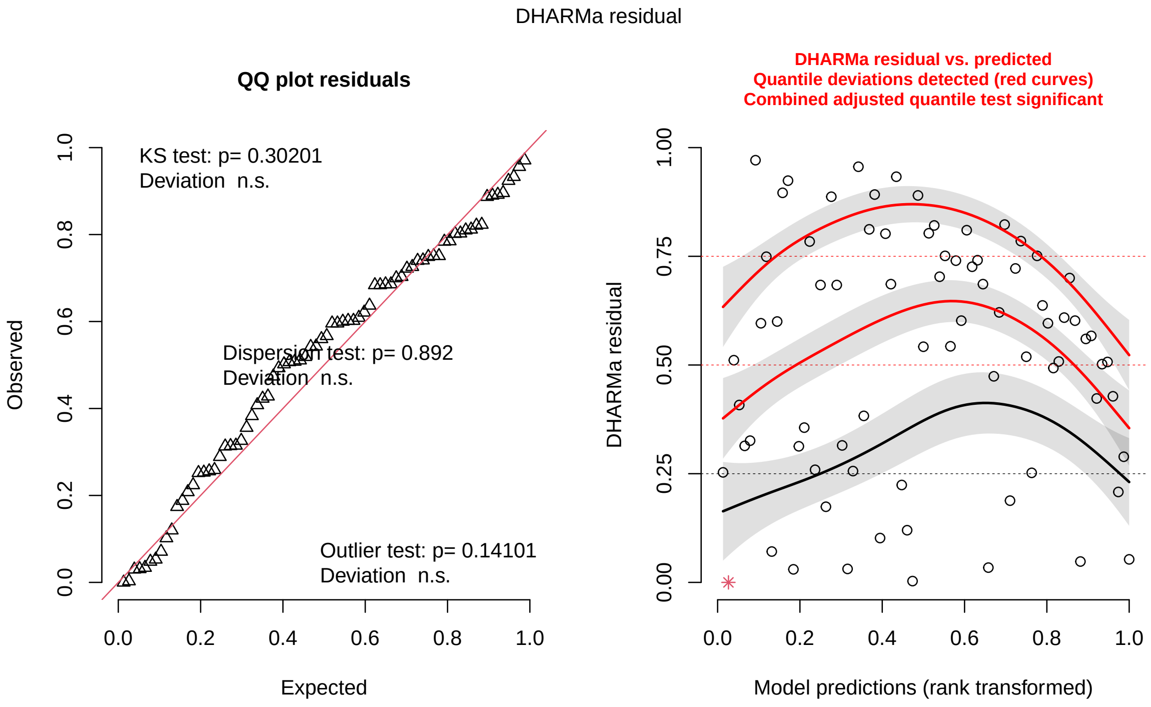

```
testDispersion(sim_residuals_auc)
```

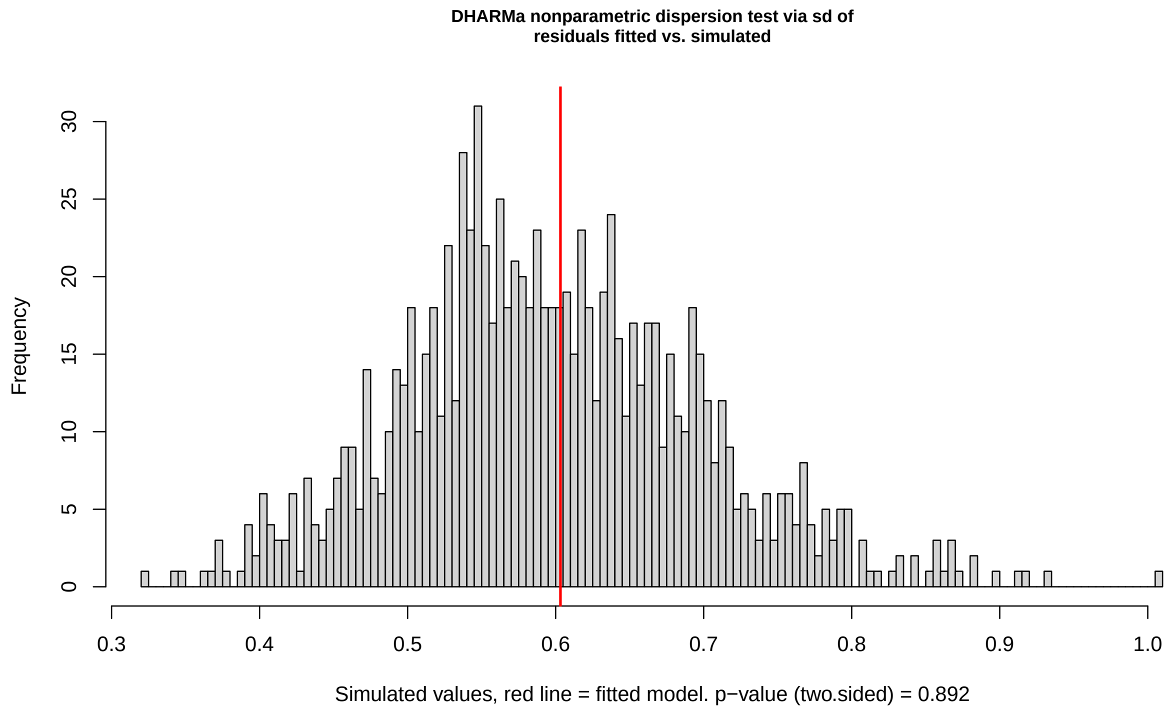

```
    DHARMa nonparametric dispersion test via sd of residuals
    fitted vs. simulated

data:  simulationOutput
dispersion = 1, p-value = 0.9
alternative hypothesis: two.sided
```

##### 3.5 Sex Post-hoc analysis for Tolerance: The correlation between the artificial selection line and tissue specific tolerance "ppmass" is being assessed for significant differences from 0 in either sex.

```
### for the females and males
FBCModel <- glmmTMB(ppmass ~ line + respreL + gen + dose + JulianDate,
    data = female, family = gaussian(link = "identity"), na.action = na.omit)

# Stats for Model
Anova(FBCModel)
```

```
Analysis of Deviance Table (Type II Wald chisquare tests)

Response: ppmass
           Chisq Df Pr(>Chisq)
line       23.81  2    6.7e-06 ***
respreL     6.96  1     0.0083 **
gen         0.21  3     0.9755
dose        0.00  1     0.9723
```

```
JulianDate  2.84  4     0.5857
---
Signif. codes:  0 '***' 0.001 '**' 0.01 '*' 0.05 '.' 0.1 ' ' 1
```

```r
AIC(FBCModel)
```

```
[1] 255
```

```r
summary(glht(FBCModel, linfct = mcp(line = "Tukey")))
```

```
     Simultaneous Tests for General Linear Hypotheses

Multiple Comparisons of Means: Tukey Contrasts

Fit: glmmTMB(formula = ppmass ~ line + respreL + gen + dose + JulianDate,
    data = female, family = gaussian(link = "identity"), na.action = na.omit,
    ziformula = ~0, dispformula = ~1)

Linear Hypotheses:
           Estimate Std. Error z value Pr(>|z|)
r - c == 0    -7.04       1.80   -3.92  0.00029 ***
s - c == 0     1.49       2.27    0.66  0.78662
s - r == 0     8.54       2.11    4.05  0.00015 ***
---
Signif. codes:  0 '***' 0.001 '**' 0.01 '*' 0.05 '.' 0.1 ' ' 1
(Adjusted p values reported -- single-step method)
```

```r
# visualization of results and verifying model with residuals
# Diagnostic plots
sim_residuals_auc <- simulateResiduals(FBCModel, 1000)
plot(sim_residuals_auc)
```

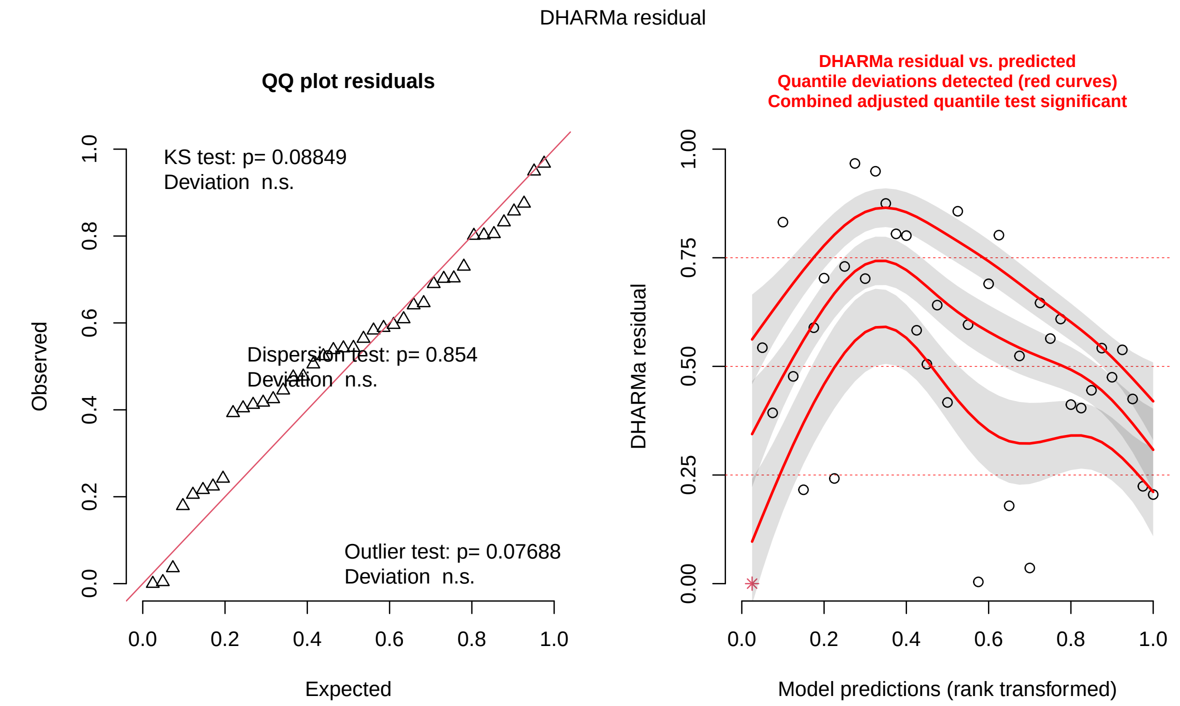

```
testDispersion(sim_residuals_auc)
```

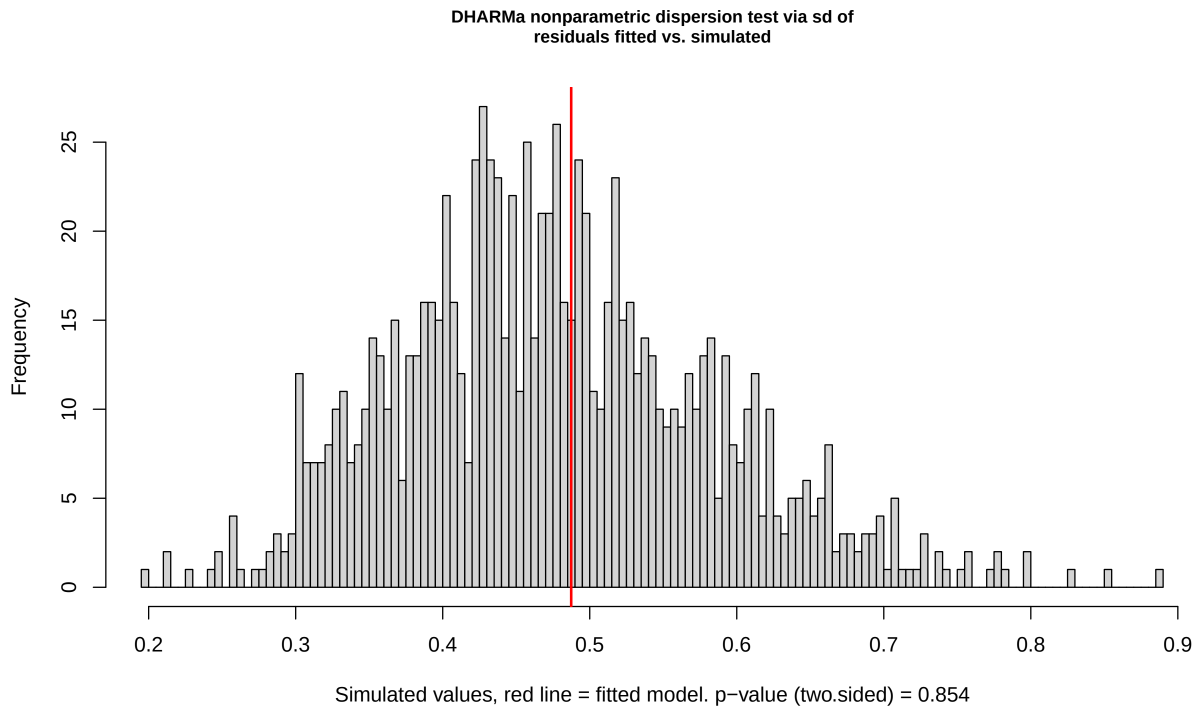

```
DHARMa nonparametric dispersion test via sd of residuals
```

```
    fitted vs. simulated

data:  simulationOutput
dispersion = 1, p-value = 0.9
alternative hypothesis: two.sided
```

```
#### males

## Model
MBCModel <- glmmTMB(ppmass ~ line + respreL + gen + dose + JulianDate,
    data = male, family = gaussian(link = "identity"), na.action = na.omit)

# Stats for Model
Anova(MBCModel)
```

```
Analysis of Deviance Table (Type II Wald chisquare tests)

Response: ppmass
           Chisq Df Pr(>Chisq)
line        2.08  2       0.35
respreL     0.00  1       0.97
gen         4.41  3       0.22
dose        7.89  5       0.16
JulianDate  3.86  4       0.42
```

```
AIC(MBCModel)
```

```
[1] 249.2
```

```
summary(glht(MBCModel, linfct = mcp(line = "Tukey")))
```

```
     Simultaneous Tests for General Linear Hypotheses

Multiple Comparisons of Means: Tukey Contrasts

Fit: glmmTMB(formula = ppmass ~ line + respreL + gen + dose + JulianDate,
    data = male, family = gaussian(link = "identity"), na.action = na.omit,
    ziformula = ~0, dispformula = ~1)

Linear Hypotheses:
           Estimate Std. Error z value Pr(>|z|)
r - c == 0     1.68       2.50    0.67     0.78
s - c == 0     4.76       3.34    1.43     0.32
s - r == 0     3.08       2.75    1.12     0.50
(Adjusted p values reported -- single-step method)
```

```
# visualization of results and verifying model with residuals
# Diagnostic plots
sim_residuals_auc <- simulateResiduals(MBCModel, 1000)
plot(sim_residuals_auc)
```

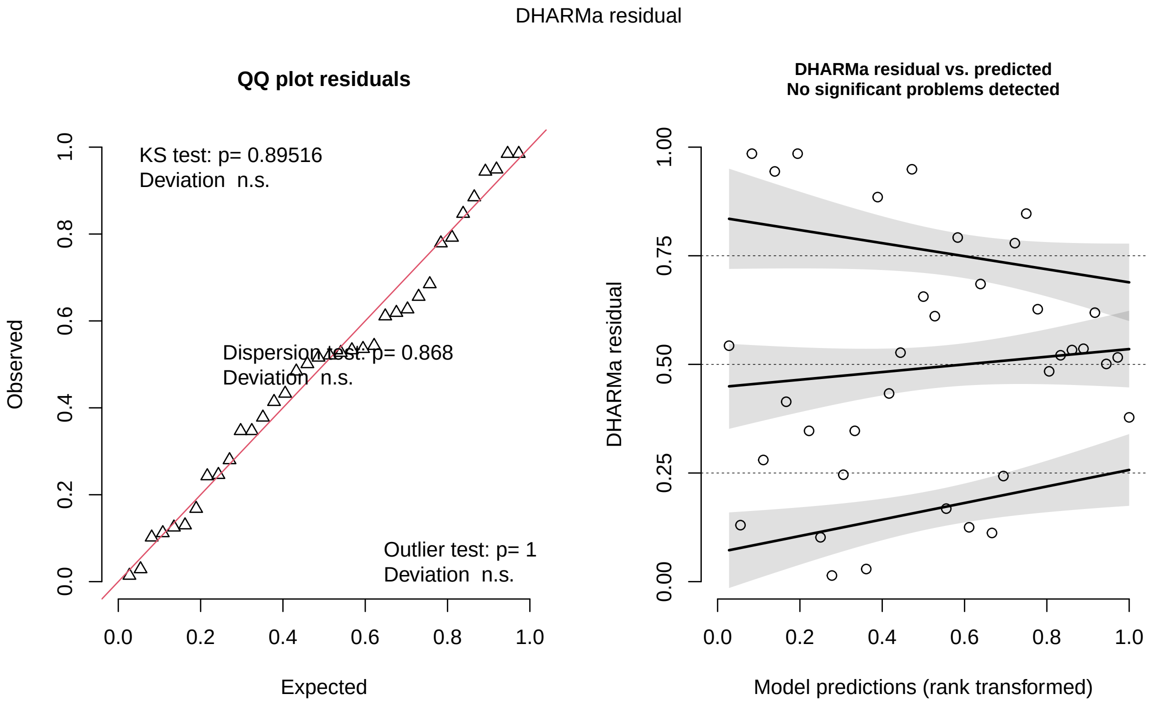

```
testDispersion(sim_residuals_auc)
```

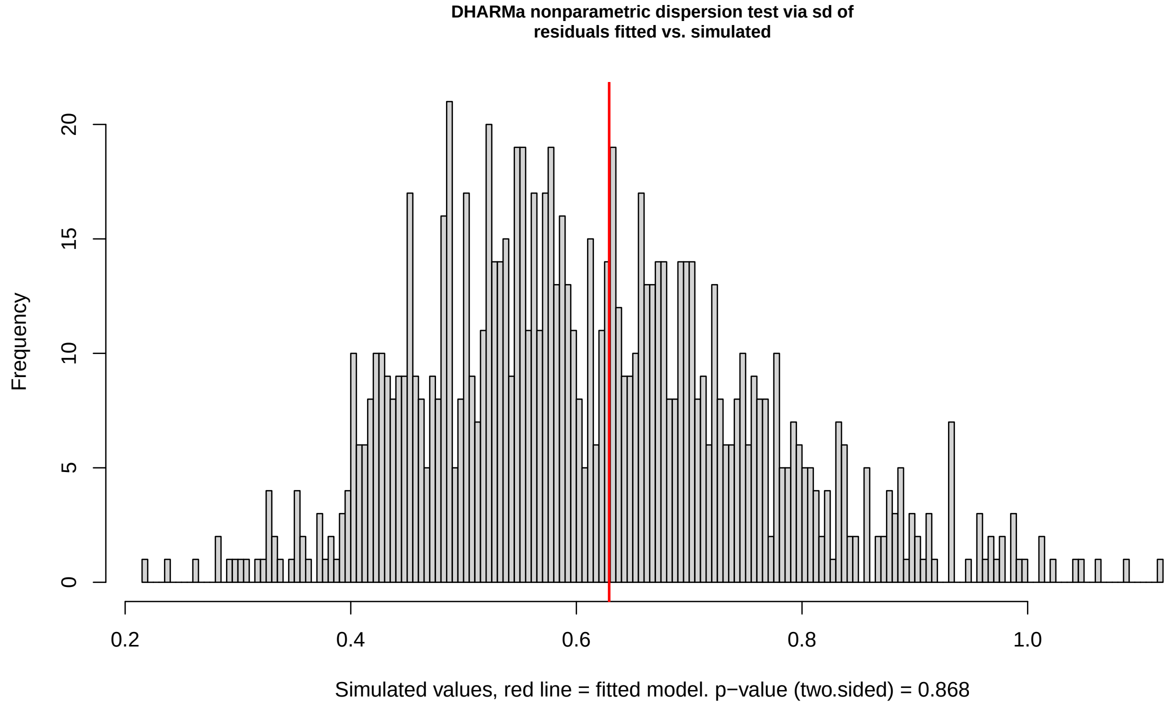

```
DHARMa nonparametric dispersion test via sd of residuals
```

```
    fitted vs. simulated

data:  simulationOutput
dispersion = 1, p-value = 0.9
alternative hypothesis: two.sided
```

##### 3.6 Evaluating the resistance-tolerance trade-off at the level of individual fish

```r
IOm$logAUC <- log(IOm$AUC)

Tradeoff_poly <- lm(ppmass ~ poly(logAUC, 2)*sex+ respreL + gen, data = IOm, na.action = na.omit)
summary(Tradeoff_poly)
```

```
Call:
lm(formula = ppmass ~ poly(logAUC, 2) * sex + respreL + gen,
    data = IOm, na.action = na.omit)

Residuals:
    Min      1Q  Median      3Q     Max
-16.349  -0.885   0.182   1.572  10.452

Coefficients:
                      Estimate Std. Error t value Pr(>|t|)
(Intercept)             -7.430      2.824   -2.63    0.011 *
poly(logAUC, 2)1        46.171      6.563    7.04  1.4e-09 ***
poly(logAUC, 2)2        -9.444      6.082   -1.55    0.125
sexm                     0.878      1.228    0.71    0.477
respreL                  0.194      0.242    0.80    0.426
gen4                     1.057      2.862    0.37    0.713
gen5                     0.729      2.936    0.25    0.805
gen6                    -0.218      2.982   -0.07    0.942
poly(logAUC, 2)1:sexm    8.870     14.277    0.62    0.537
poly(logAUC, 2)2:sexm   -5.282     16.970   -0.31    0.757
---
Signif. codes:  0 '***' 0.001 '**' 0.01 '*' 0.05 '.' 0.1 ' ' 1

Residual standard error: 3.91 on 66 degrees of freedom
  (11 observations deleted due to missingness)
Multiple R-squared:  0.666, Adjusted R-squared:  0.62
F-statistic: 14.6 on 9 and 66 DF,  p-value: 1.11e-12
```

```r
Anova(Tradeoff_poly)
```

```
Anova Table (Type II tests)

Response: ppmass
                    Sum Sq Df F value  Pr(>F)
poly(logAUC, 2)       1698  2   55.49 7.3e-15 ***
sex                     10  1    0.65    0.42
respreL                 10  1    0.64    0.43
```

```
gen                        17  3    0.37    0.77
poly(logAUC, 2):sex        15  2    0.49    0.61
Residuals                1010 66
---
Signif. codes:  0 '***' 0.001 '**' 0.01 '*' 0.05 '.' 0.1 ' ' 1
```

```r
##Visualize risduals
p <- visreg(Tradeoff_poly, "logAUC", gg = TRUE) +
  labs(x = "Infection Severity (worm days)", y = "Relative Tissue-Specific Tolerance") +  # Custom axis
  theme_minimal() +
  theme(
    axis.title = element_text(size = 16),  # Increase font size for axis titles
    axis.text = element_text(size = 14)    # Increase font size for axis text
  ) +
  geom_text(aes(x = mean(IOm$logAUC), y = max(IOm$ppmass),  # Position of the slope text
              label = paste("Slope:", round(coef(Tradeoff_poly)[2], 3))),
            size = 5, color = "blue")
```

```
Conditions used in construction of plot
sex: f
respreL: -0.2158
gen: 4
```

#### 3.7 Genetic Analysis: Investigation of whether results observed can be attributed to the effects of inbreeding within the selection lines

We evaluated the genetic diversity and variance in inbreeding within the wild-caught populations, F1, F3-F7 generations in order to determine if the observed difference in response to infection between the resistant and susceptible lines was due potentially induced by reduced population sizes. We are showing one example of the code ran for all populations. **For all data see supplemental tables 1-5.** #Identity disequilibrium is not significant in inbreeding nor is there a different in the amount of genetic diversity (seen in supplement table below). ). To get FIS values were assessed in GENEPOP ((web version 4.2; Raymond and Rousset 1995).

Previous authors have demonstrated that inbred Poecilia reticulata individuals infected with Gyrodactylus turnbulli exhibited significantly higher mean intensity of infection (Smallbone et al. 2016). We evaluated the genetic diversity and variance in inbreeding within the wild-caught populations, F1, F3-F7 generations in order to determine if the observed difference in response to infection between the resistant and susceptible lines was due potentially induced by reduced population sizes. DNA was isolated from fin clips with the HotSHOT method by Truett, et al. (2000) using 50 µl of alkaline lysis reagent and neutralizing solution followed by incubation at 95 °C for 30 minutes. Eight previously designed microsatellite markers were amplified in two multiplex assays. Multiplex assay one included Pret 77 and Pret 69 (Watanabe et al. 2003), PP-GATA-5 (Nater et al. 2008), and Pre9 (Patterson et al. 2005). Multiplex assay 2 included Hull 9-1 (van Oosterhout, et al. 2006), Pre92 (Becher et al. 2002), Pre15 and Pre 18 (Paterson et al. 2005). Forward primers were fluorescently labelled and reverse primers had a short tail sequence (GTTTCCT) added at the 5'-end to reduce polyadenylation in all loci except Pre9, as the modification prevented amplification at this locu (Brownstein, et al. 1996). Multiplex assays contained 10 µl Type-It Master Mix (Qiagen, Hilden, Germany), 2 µl of multiplex primer mixes (Assay 1: 0.2 mM each for forward and reverse primer for Pre 77, 0.5 mM each for forward and reverse primer for PP-GATA-5, 0.1 mM each for forward and reverse primer for Pret 69, and 0.2 mM each for forward and reverse primer for Pret 9; Assay 2: 0.4 mM each for forward and reverse primer for Hull 9-1, 0.05 mM each for forward and reverse primer for Pre 92, 0.5 mM each for forward and reverse primer for Pre 15, and 0.05 mM each for forward and reverse primer for Pre 18), 2 µl Q-solution (Qiagen, Hilden, Germany) 4 µl molecular grade $H_2O$ and 2 µl template DNA. PCR reaction conditions included an initial denaturation at 95 °C for 5 minutes, followed by 35 cycles of denaturation for 30 seconds at 94 °C, annealing for 90 seconds at 60 °C or 56 °C (for MP1 and MP2, respectively) and elongation for 30 seconds at 72 °C. After a final elongation step for 30 minutes at 60 °C, PCR products were kept frozen until further processing. The PCR product was visualized on a X % agarose gel, stained with ethidium bromide (insert concentration here) and observed under ultraviolet light. Samples with visible PCR product were genotyped on an ABI Genetic Analyzer 3730 (Applied Biosystems, USA). Samples were scored using GeneMarker (Version 2.6.4; Softgenetics) and binned using the package MSatAllele (Alberto 2013) in R (R Core Team 2018).

CF_data.txt: subset of control female fish microstats data

Here is the example code:

```
invisible({
    capture.output({

        # g2 is identity disequilibrium Tells you the correlation of
        # across multi-locus phenotype, so you want a low number here

        CF_data = read.table(file = "CF_data.txt", sep = "\t", header = TRUE)

        guppy_microsats <- convert_raw(CF_data)

        g2_guppy_microsats <- g2_microsats(guppy_microsats, nperm = 999,
            nboot = 100, CI = 0.95)

    })
})
```

```
options(width = 70, digits = 4)
# Printing results
print(g2_guppy_microsats)
```

```
Calculation of identity disequilibrium with g2 for microsatellite data
----------------------------------------------------------------------

Data: 21 observations at 8 markers
Function call = g2_microsats(genotypes = guppy_microsats, nperm = 999, nboot = 100,     CI = 0.95)

g2 = -0.04292, se = 0.03834

confidence interval
   2.5%   97.5%
-0.1168  0.0110

p (g2 > 0) = 0.7828 (based on 999 permutations)
```

#### 3.8 Calculating Hertibility in artifical selection lines from Breeders equation

For the heritibility analysis we used “ParentalSelection2.csv” to find the mean of meanworm and tolerance of the founders of each line. We rescaled both “meanworm” and “ppmass” due to the differences in parasite growth from the first round of selection to the infection of offspring. This allowed us to have comparable means. Figures show directly founders and offspring have responded to selection in their meanworm and tolerance.

```
## founders of the lines

s1 <- subset(Parental, line == "s1")  #subset parental phenotype susceptible

mean(s1$rescalep)  # 20.54 this is the average selected meanworm phenotype for Susceptible parents
```

```
[1] NA
```

```
# centered mean
mean(s1$centermw)  #8.981
```

```
[1] 8.981
```

```
s1b <- s1[!is.na(s1$ppmass), ]
mean(s1b$centerpp)  #5.555
```

```
[1] 5.555
```

```
r1 <- subset(Parental, line == "r1")  #subset parental phenotype resistant
mean(r1$centermw)  #-8.917 this is the average selected meanworm phenotype for Resistant parents
```

```
[1] -8.917
```

```
r1b <- r1[!is.na(r1$ppmass), ]
mean(r1b$centerpp)  #-12.1
```

```
[1] -12.1
```

```
## Offspring

so <- subset(IO, line == "s")
mean(so$centermw)  #7.057 this is the average centered response phenotype for S
```

```
[1] 7.057
```

```
sob <- so[!is.na(so$ppmass), ]
mean(sob$centerpp)  #2.918
```

```
[1] 2.918
```

```
ro <- subset(IO, line == "r")
mean(ro$centermw)  #-3.662
```

```
[1] -3.662
```

```r
rob <- ro[!is.na(ro$ppmass), ]
mean(rob$centerpp)  #-2.972
```

```
[1] -2.972
```

```r
## Calculating heritability for Susceptible lines meanworm

((7.057 - 0)/(8.981 - 0))
```

```
[1] 0.7858
```

```r
# this equals 0.7858

## Calculating heritability for Susceptible lines tolerance

((2.918 - 0)/(5.555 - 0))
```

```
[1] 0.5253
```

```r
# this equals 0.5253

## Calculating heritability for Resistant lines meanworm
((-3.662 - 0)/(-8.917 - 0))
```

```
[1] 0.4107
```

```r
# this equals 0.4107

## Calculating heritability for Resistant lines tolerance
((-2.87 - 0)/(-12.1 - 0))
```

```
[1] 0.2372
```

```r
# this equals 0.2372
```

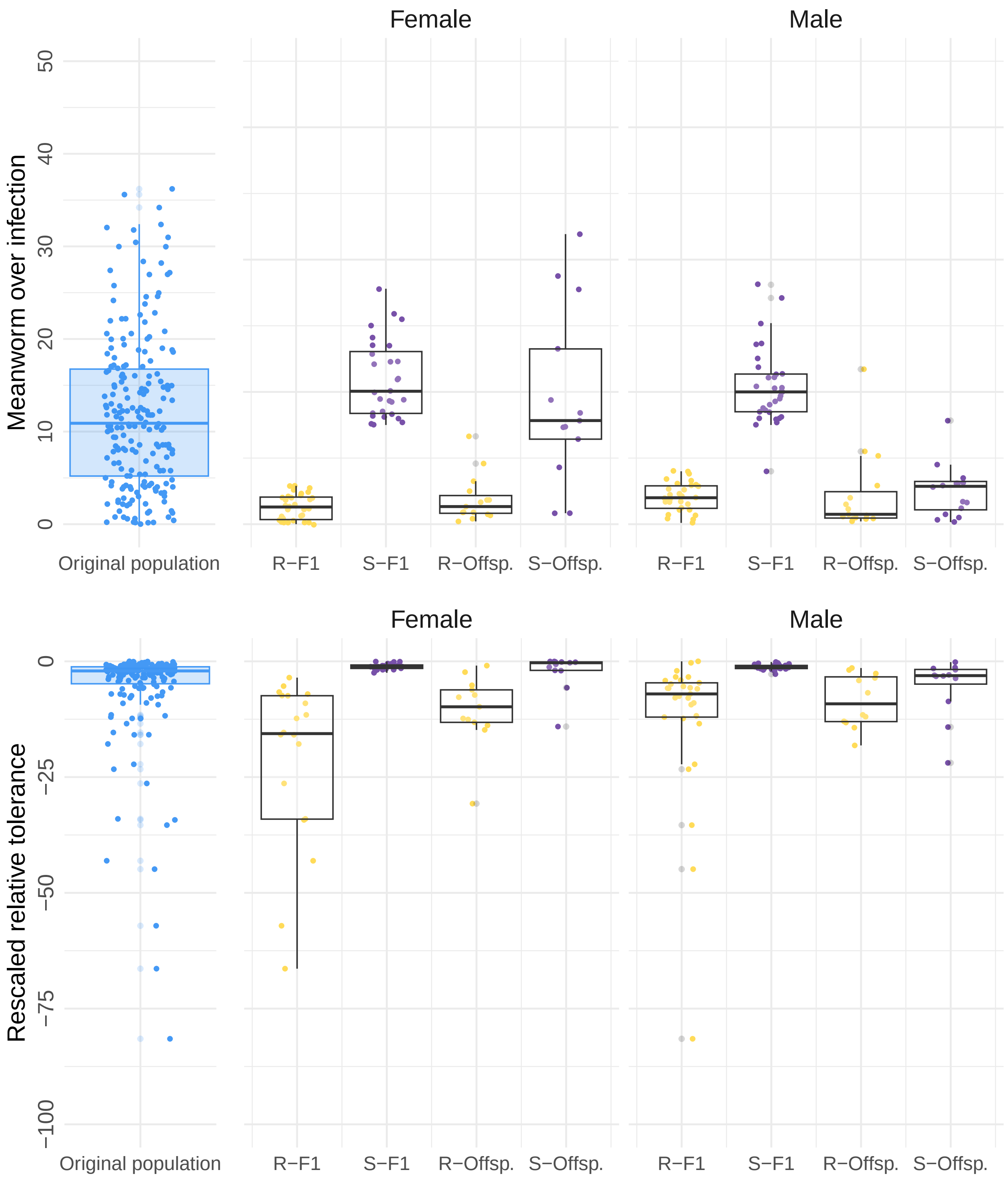

Female
Male
Meanworm over infection
50
40
30
20
10
0
Original population
R−F1
S−F1
R−Offsp.
S−Offsp.
R−F1
S−F1
R−Offsp.
S−Offsp.
Female
Male
Rescaled relative tolerance
0
−25
−50
−75
−100
Original population
R−F1
S−F1
R−Offsp.
S−Offsp.
R−F1
S−F1
R−Offsp.
S−Offsp.

### 4 Part II: Looking deeper:The host-associated microbiome clustering in artifical selection lines or independently coorelating with infections integral

There is evidence that the tiny organisms living in and on hosts called the microbiome, might be passed down from one generation to the next and have evolved alongside their hosts (a conponent of the "holobiont" theory). To find out whether the microbiome on the skin of our fish played a role in making them more or less resistant to infections, we collected samples from 51 fish: Control (n=15), Resistant (n=18), Susceptible (n=17), Sham infected fish (n=20). We then analyzed the communities of microorganisms in these samples using 16s rRNA sequencing. Our goal was to see if the fish from different lines had different associated skin microbiomes, and if these microbiomes were related to how susceptible or resistant they were to infections.

In continuation of the above analysis, the most important data can be found in the file 'SelectionData.csv,' This section expands to microbial community dataframes:

- **Unweighted-distance-matrix.tsv and Weighted-distance-matrix.tsv**: Beta-diversity obtained from Qiime2 export pipeline- Unweighted and Weighted Unifrac distance matrices for PCoA creation and PERMANOVA analysis to which factor explain microbial variation on guppies skin.
- **Discrete__abundances.csv**: Relative frequencies of bacterial taxa data table exported from Qiime2 after DADA2 processing and SILVA classification.
- **pred__metagenome__unstrat.tsv:** PiCrust2 predicted KO (KEGG Orthology) abundances normalized by 16s copy number and converted to relative abundance for visualization

**Sub-setting of SelectionData.csv explain:**

**M1 Data frame**: Subset (n=51) F4-7 offspring fish (sham infected included) that were swabbed and sent for 16s rRNA 515F–806R sequencing.

**MicroDat Data frame**: Subset (n=31) of parasite infected offspring fish that were swabbed and sent for 16s rRNA 515F–806R sequencing.

#### 4.1 Artifical selections lines significance in explaining alpha diversity or beta diversity

**Alpha Diversity** of each line (C,R,S)was tested using Pairwise Kruskal-Wallis preformed on QIIME2 for Shannon's index, Faith's Phylogenic Diversity (PD), and ASVs.

- *Shannon's Index*: C (n=15), R (n=18), S (n=17) H=0.16 q-value=0.92
- *Faith's PD*: C (n=15), R (n=18), S (n=17) H=0.028 q-value=0.98
- *ASVs*: C (n=15), R (n=18), S (n=17) H=1.74 q-value=0.41

```
## Do C,R,S lines explains a significant amount of variance in beta
## diversity?
options(width = 70, digits = 4)
## PERMANOVA
set.seed(1)
UnweightResults <- adonis2(formula = UNW_matrix ~ line + sex + respreL +
    rPREsmi, data = UnWeighted, by = "margin", permutations = 999)
print(UnweightResults)
```

```
Permutation test for adonis under reduced model
Marginal effects of terms
Permutation: free
Number of permutations: 999

adonis2(formula = UNW_matrix ~ line + sex + respreL + rPREsmi, data = UnWeighted, permutations = 999, by
         Df SumOfSqs    R2    F Pr(>F)
line      2     0.35 0.033 0.81  0.955
sex       1     0.33 0.032 1.56  0.006 **
respreL   1     0.21 0.021 1.01  0.429
rPREsmi   1     0.20 0.019 0.95  0.577
Residual 44     9.36 0.898
Total    49    10.42 1.000
---
Signif. codes:  0 '***' 0.001 '**' 0.01 '*' 0.05 '.' 0.1 ' ' 1
```

##### 4.2 Due the different sex composition result with Unweighted UniFrac, tested Alpha diversity differences due to fish sex

**Alpha Diversity** of Fish Sex Pairwise Kruskal-Wallis preformed on QIIME2 for Shannon's index and ASVs

- *Shannon's Index*: f (n=23) m (n=27) H=4.0588245 q-value=0.043941
- *Faith's PD*: f (n=23) m (n=27) H=4.71 q-value=0.030
- *ASVs*: f (n=23) m (n=27) H=4.883137 q-value=0.02712

##### 4.3 DMM Modeling used to parse out bacterial communties types.

We did not see any effect of microbial clustering within the artificial selection lines. To see if there is any patterns or clusters within our fish's microbial communities we used a technique developed for finding communities types based on species type and abundance of bacteria present in microbial communities on each fish. We then can see if any communities cluster into artificial selection.

**Dirichlet multinomial mixture model**: (Quince et al. 2012) is a probabilistic method for community typing (or clustering) of microbial community profiling data. It is an infinite mixture model, which means that the method can infer the optimal number of community types.

#Laplace Appromation found three communities fit data best

```r
## Setting up the display parameters and data upload set width of R
## output to 70 characters and number of floating point digits
## displayed to two
options(width = 70, digits = 4)

# use .qualitative color set
.qualitative <- DirichletMultinomial:::.qualitative

# dev.off redefined to return without displaying values dev.off <-
# function(...) invisible(grDevices::dev.off(...))

## distribution of reads from each taxon, on a log scale (figure 1)
cnts <- log10(colSums(DMMdata))
plot(densityplot(cnts, xlim = range(cnts), xlab = "Taxon representation (count)"))
```

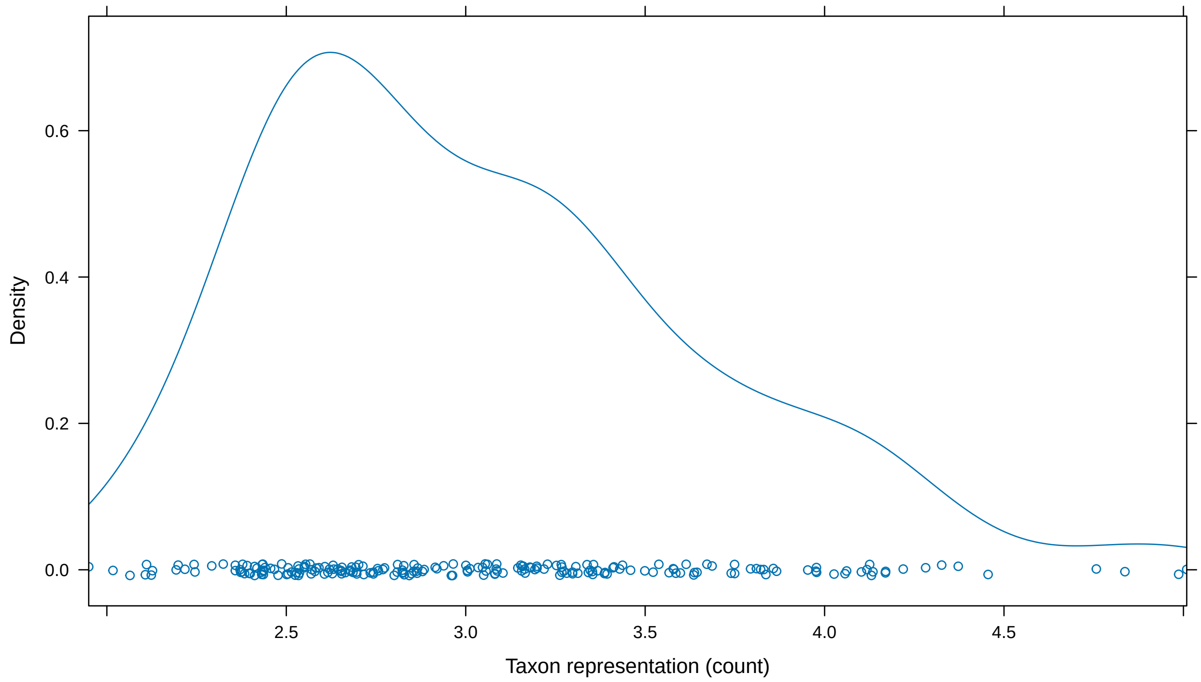

```
# DMM Part2 Finding number of clusters Identifying number of
# community types in study population fit count data to a series of
# models with 1 to 8 community types important: the higher the
# maximum number of community types, the higher the computing time
# (exponentially increases) I've set the maximum number to 8 due to
# computational time and because the line is continually increasing
# You may want to consider going up to at least 10 if you are trying
# this with your own data - WARNING: will take you all day unless you
# have a supercomputer

# fit <- mclapply(1:8, dmn, count=DMMdata, verbose=TRUE)

# Saved for later so you don't have to run each time
load("fit.rda")
```

```
### The return value can be queried for measures of fit (Laplace,
### AIC, BIC); these are plotted for different k

lplc <- sapply(fit, laplace)

plot(lplc, type = "b", xlab = "Number of Dirichlet Components", ylab = "Model Fit")
```

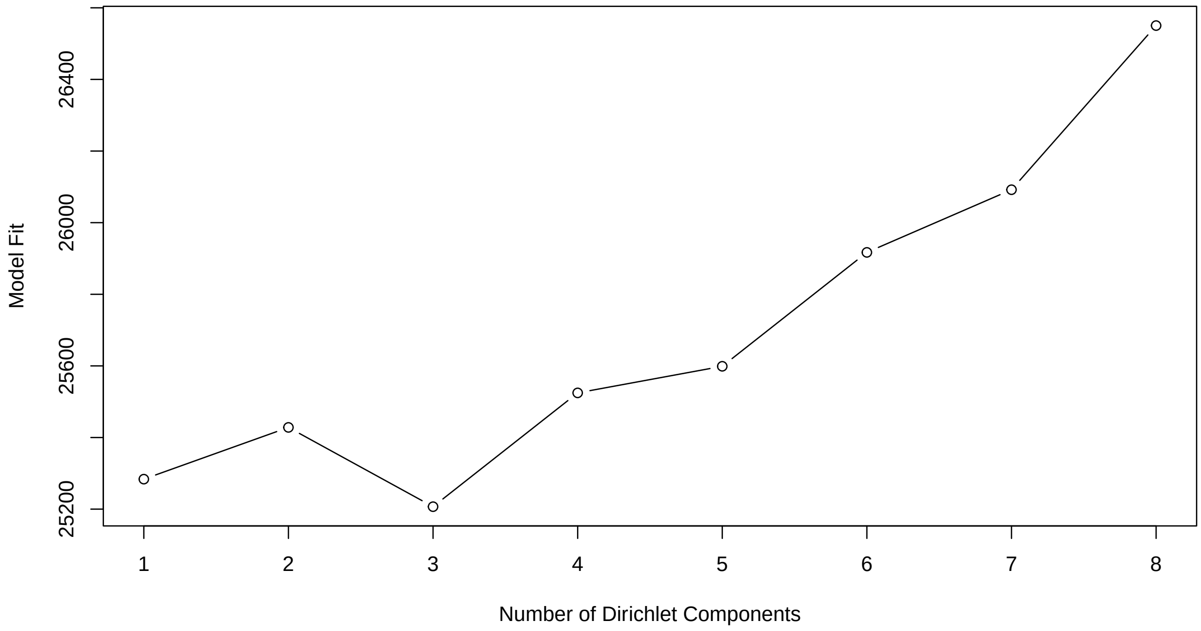

```
###### Characterizing best model pull out best model and save as
###### 'best'

(best <- fit[[3]])
```

```
class: DMN
k: 3
samples x taxa: 50 x 214
Laplace: 25207 BIC: 26862 AIC: 26246
```

```
### plot contribution of each taxonomic group to community types

plot(splom(log(fitted(best))))
```

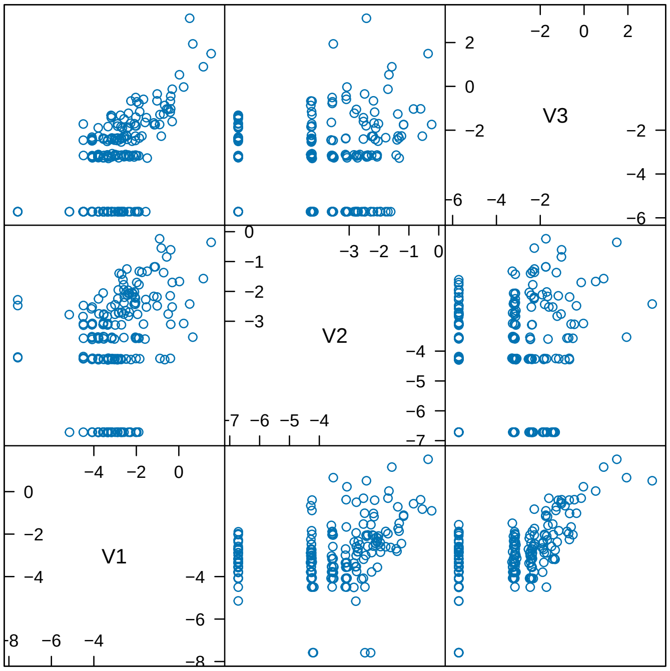

Scatter Plot Matrix

#### 4.4 Does Bacterial community have a uniform distribution across line?

The assumed null hypothesis states that the Bacterial_community is evenly distributed across the line. Since the p-value is much larger than the typical significance level (e.g., 0.05), you can conclude that there is no statistically significant difference in the distribution of bacterial communities between the artificial selection lines. Therefore, bacterial communities are evenly distributed across the artifical selection lines.

```
# Create contingency table for line and bacterial community
contingency_table <- table(M1$Bacterial_Community, M1$line)
# Perform Fisher's Exact Test
fisher_test <- fisher.test(contingency_table)
# View results
fisher_test
```

```
    Fisher's Exact Test for Count Data

data:  contingency_table
p-value = 0.8
alternative hypothesis: two.sided
```

#Checking that the bacterial communities types found significantly explain variation in bacterial community structure

```
options(width = 70, digits = 4)

## PERMANOVA
WeightResults <- adonis2(formula = W_matrix ~ Bacterial_Community + sex +
```

```r
    respreL + rPREsmi, data = Weighted, by = "margin", permutations = 999)
print(WeightResults)
```

```
Permutation test for adonis under reduced model
Marginal effects of terms
Permutation: free
Number of permutations: 999

adonis2(formula = W_matrix ~ Bacterial_Community + sex + respreL + rPREsmi, data = Weighted, permutation
                    Df SumOfSqs    R2    F Pr(>F)
Bacterial_Community  2    0.515 0.247 7.85  0.001 ***
sex                  1    0.032 0.015 0.97  0.430
respreL              1    0.021 0.010 0.64  0.773
rPREsmi              1    0.044 0.021 1.36  0.179
Residual            44    1.443 0.691
Total               49    2.087 1.000
---
Signif. codes:  0 '***' 0.001 '**' 0.01 '*' 0.05 '.' 0.1 ' ' 1
```

```r
UnweightResults <- adonis2(formula = UNW_matrix ~ Bacterial_Community +
    sex + respreL + rPREsmi, data = UnWeighted, by = "margin", permutations = 999)
print(UnweightResults)
```

```
Permutation test for adonis under reduced model
Marginal effects of terms
Permutation: free
Number of permutations: 999

adonis2(formula = UNW_matrix ~ Bacterial_Community + sex + respreL + rPREsmi, data = UnWeighted, permuta
                    Df SumOfSqs    R2    F Pr(>F)
Bacterial_Community  2     0.91 0.087 2.27  0.001 ***
sex                  1     0.26 0.025 1.29  0.068 .
respreL              1     0.20 0.019 0.99  0.473
rPREsmi              1     0.20 0.019 0.98  0.477
Residual            44     8.80 0.844
Total               49    10.42 1.000
---
Signif. codes:  0 '***' 0.001 '**' 0.01 '*' 0.05 '.' 0.1 ' ' 1
```

```r
## The communities found do significantly explain variance found in
## bacterial composition
```

#### 4.5 Visualization of data with Bacterial community subset

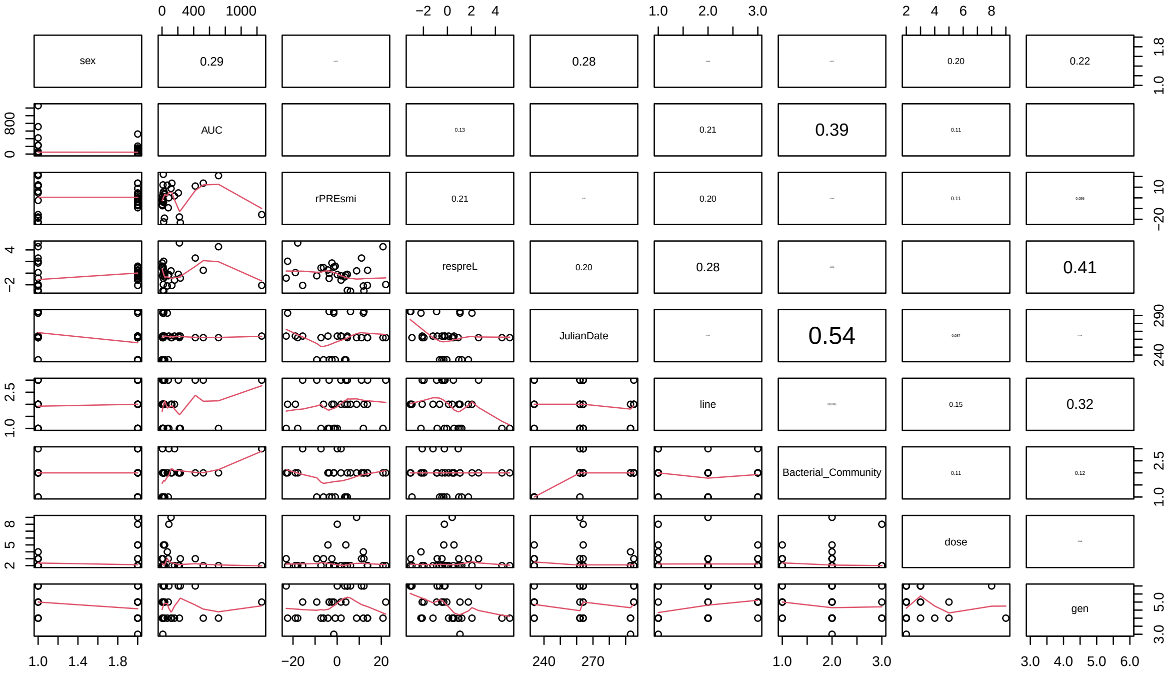

#### 4.6 GLMM: Bacterial community type explains variation in *Gyrodactylus turnbulli* infection integral.

**Response variable:** the area under the curve for worm days/ time "infection integral".

**Fixed Effects:**

- *Bacterial_Community*: Community state type of the microbiome assigned by DMM model
- *Line* (F4-7 Resistant, susceptible, control fish)
- *rPREsmi* (residuals of scaled mass index "body condition" of fish)
- *respreL* (residual of fish lengths)
- *Sex* of fish
- *Julian Date* (Date fish were infected)

```r
options(width = 70, digits = 4)
## Microbiome model
BacLinemod <- glmmTMB(AUC ~ Bacterial_Community * line + respreL + rPREsmi +
    sex * line + JulianDate + gen + dose, data = MicroDat, family = Gamma(link = "log"))
AIC(BacLinemod)
```

```
[1] 342.2
```

```r
summary(BacLinemod)
```

```
 Family: Gamma  ( log )
Formula:
AUC ~ Bacterial_Community * line + respreL + rPREsmi + sex *
    line + JulianDate + gen + dose
Data: MicroDat

     AIC      BIC   logLik deviance df.resid
   342.2    368.0   -153.1    306.2       13

Dispersion estimate for Gamma family (sigma^2): 0.614

Conditional model:
                           Estimate Std. Error z value Pr(>|z|)
(Intercept)                 12.0016     4.4768    2.68   0.0073 **
Bacterial_Community1        -2.2283     0.5052   -4.41    1e-05 ***
Bacterial_Community2         0.3593     0.3844    0.93   0.3499
line1                       -0.5735     0.3231   -1.78   0.0759 .
line2                       -0.3678     0.3194   -1.15   0.2496
respreL                      0.1569     0.1128    1.39   0.1641
rPREsmi                      0.0377     0.0172    2.19   0.0282 *
sex1                         0.1267     0.2373    0.53   0.5934
JulianDate                  -0.0393     0.0164   -2.39   0.0170 *
gen                          0.5120     0.2739    1.87   0.0616 .
dose                         0.0421     0.1082    0.39   0.6973
Bacterial_Community1:line1   0.7121     0.4443    1.60   0.1090
Bacterial_Community2:line1   0.7958     0.4691    1.70   0.0898 .
Bacterial_Community1:line2  -0.2628     0.3846   -0.68   0.4944
Bacterial_Community2:line2   0.2645     0.3567    0.74   0.4583
line1:sex1                   0.7112     0.3024    2.35   0.0187 *
line2:sex1                  -0.0422     0.3357   -0.13   0.8999
---
Signif. codes:  0 '***' 0.001 '**' 0.01 '*' 0.05 '.' 0.1 ' ' 1
```

```r
Anova(BacLinemod)
```

```
Analysis of Deviance Table (Type II Wald chisquare tests)

Response: AUC
                         Chisq Df Pr(>Chisq)
Bacterial_Community      19.83  2    4.9e-05 ***
line                      8.73  2      0.013 *
respreL                   1.94  1      0.164
rPREsmi                   4.82  1      0.028 *
sex                       0.65  1      0.421
JulianDate                5.70  1      0.017 *
gen                       3.49  1      0.062 .
dose                      0.15  1      0.697
Bacterial_Community:line  7.40  4      0.116
line:sex                  7.46  2      0.024 *
---
```

```
Signif. codes:  0 '***' 0.001 '**' 0.01 '*' 0.05 '.' 0.1 ' ' 1
```

```r
summary(glht(BacLinemod, linfct = mcp(Bacterial_Community = "Tukey")))
```

```
Warning in mcp2matrix(model, linfct = linfct): covariate interactions
found -- default contrast might be inappropriate

	 Simultaneous Tests for General Linear Hypotheses

Multiple Comparisons of Means: Tukey Contrasts

Fit: glmmTMB(formula = AUC ~ Bacterial_Community * line + respreL +
    rPREsmi + sex * line + JulianDate + gen + dose, data = MicroDat,
    family = Gamma(link = "log"), ziformula = ~0, dispformula = ~1)

Linear Hypotheses:
           Estimate Std. Error z value Pr(>|z|)
B - A == 0    2.588      0.800    3.23   0.0034 **
C - A == 0    4.097      0.833    4.92   <0.001 ***
C - B == 0    1.510      0.610    2.47   0.0349 *
---
Signif. codes:  0 '***' 0.001 '**' 0.01 '*' 0.05 '.' 0.1 ' ' 1
(Adjusted p values reported -- single-step method)
```

```r
## residual plots visualization of results and verifying model with
## residuals Diagnostic plots
sim_residuals_auc <- simulateResiduals(BacLinemod, 1000)
plot(sim_residuals_auc)
```

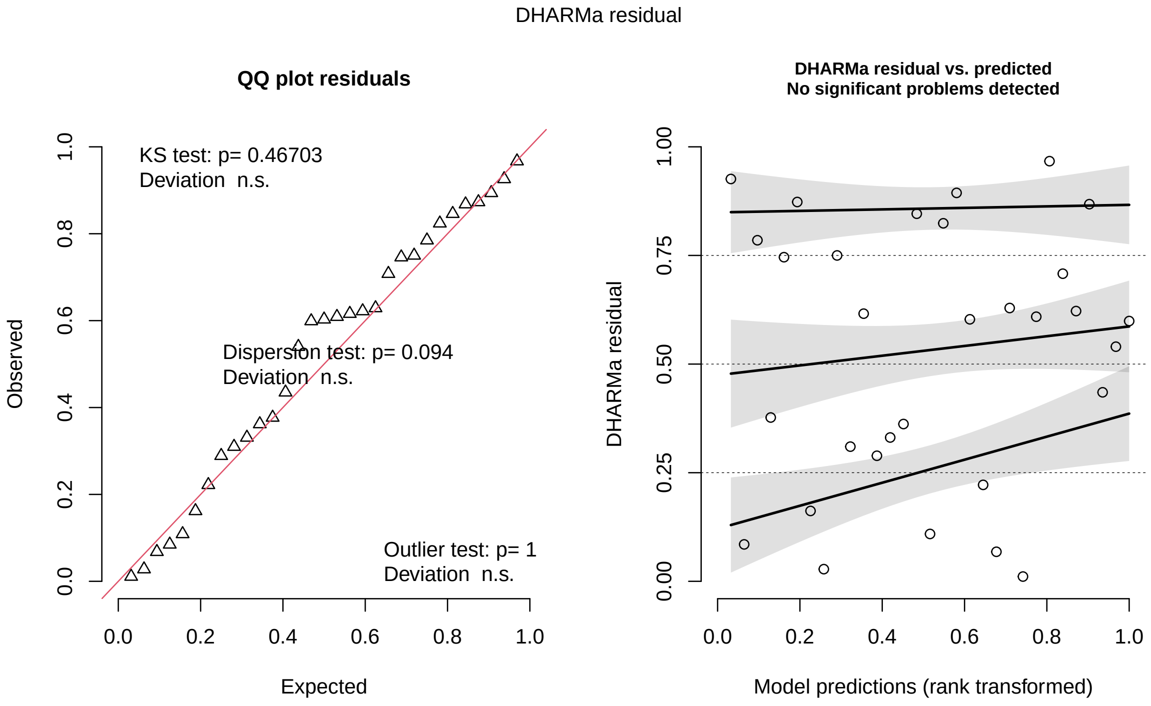

```r
testDispersion(sim_residuals_auc)
```

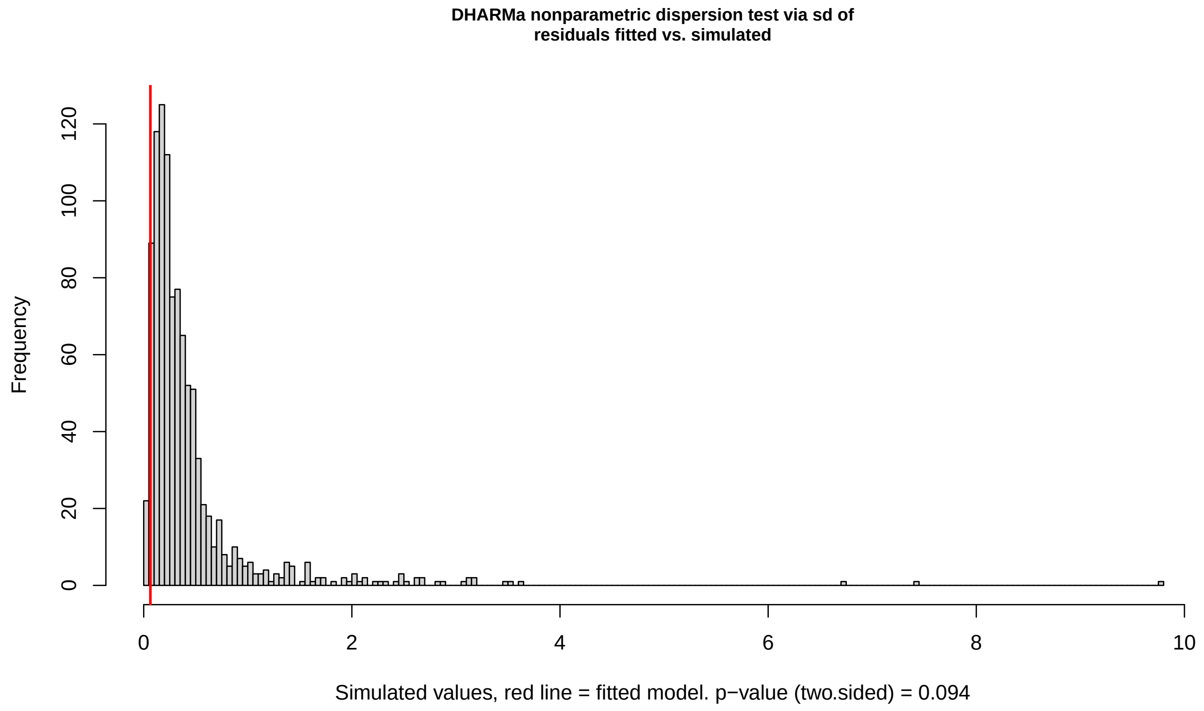

```
	DHARMa nonparametric dispersion test via sd of residuals
	fitted vs. simulated

data:  simulationOutput
dispersion = 0.15, p-value = 0.09
alternative hypothesis: two.sided
```

### 5 PART III: Investigation of explainatory power between artifical selection and microbiome.

Are the GLMM models with either the host associated microbiome or artificial selection line variables significantly better at explain infection outcomes of Trinidadian guppies?

```r
options(width = 70, digits = 4)

# Which model is significantly better at explaining infection
# severity?

LineModel <- glmmTMB(AUC ~ rPREsmi + respreL + line * sex + gen + dose +
    JulianDate + (1 | line/rep), data = MicroDat, family = Gamma(link = "log"))

LineBACModel <- glmmTMB(AUC ~ Bacterial_Community + rPREsmi + respreL +
    line * sex + gen + dose + JulianDate, data = MicroDat, family = Gamma(link = "log"))
```

```r
BACModel <- glmmTMB(AUC ~ Bacterial_Community + rPREsmi + respreL + sex +
    gen + dose + JulianDate, data = MicroDat, family = Gamma(link = "log"))

anova(LineModel, LineBACModel, BACModel)
```

```
Data: MicroDat
Models:
BACModel: AUC ~ Bacterial_Community + rPREsmi + respreL + sex + gen + dose + , zi=~0, disp=~1
BACModel:     JulianDate, zi=~0, disp=~1
LineModel: AUC ~ rPREsmi + respreL + line * sex + gen + dose + JulianDate + , zi=~0, disp=~1
LineModel:     (1 | line/rep), zi=~0, disp=~1
LineBACModel: AUC ~ Bacterial_Community + rPREsmi + respreL + line * sex + , zi=~0, disp=~1
LineBACModel:     gen + dose + JulianDate, zi=~0, disp=~1
             Df AIC BIC logLik deviance Chisq Chi Df Pr(>Chisq)
BACModel     10 347 362   -164      327
LineModel    14 354 374   -163      326  1.08      4        0.9
LineBACModel 14 341 361   -156      313 13.40      0     <2e-16 ***
---
Signif. codes:  0 '***' 0.001 '**' 0.01 '*' 0.05 '.' 0.1 ' ' 1
```

```r
AIC(LineModel, LineBACModel, BACModel)
```

```
             df   AIC
LineModel    14 354.3
LineBACModel 14 340.9
BACModel     10 347.4
```

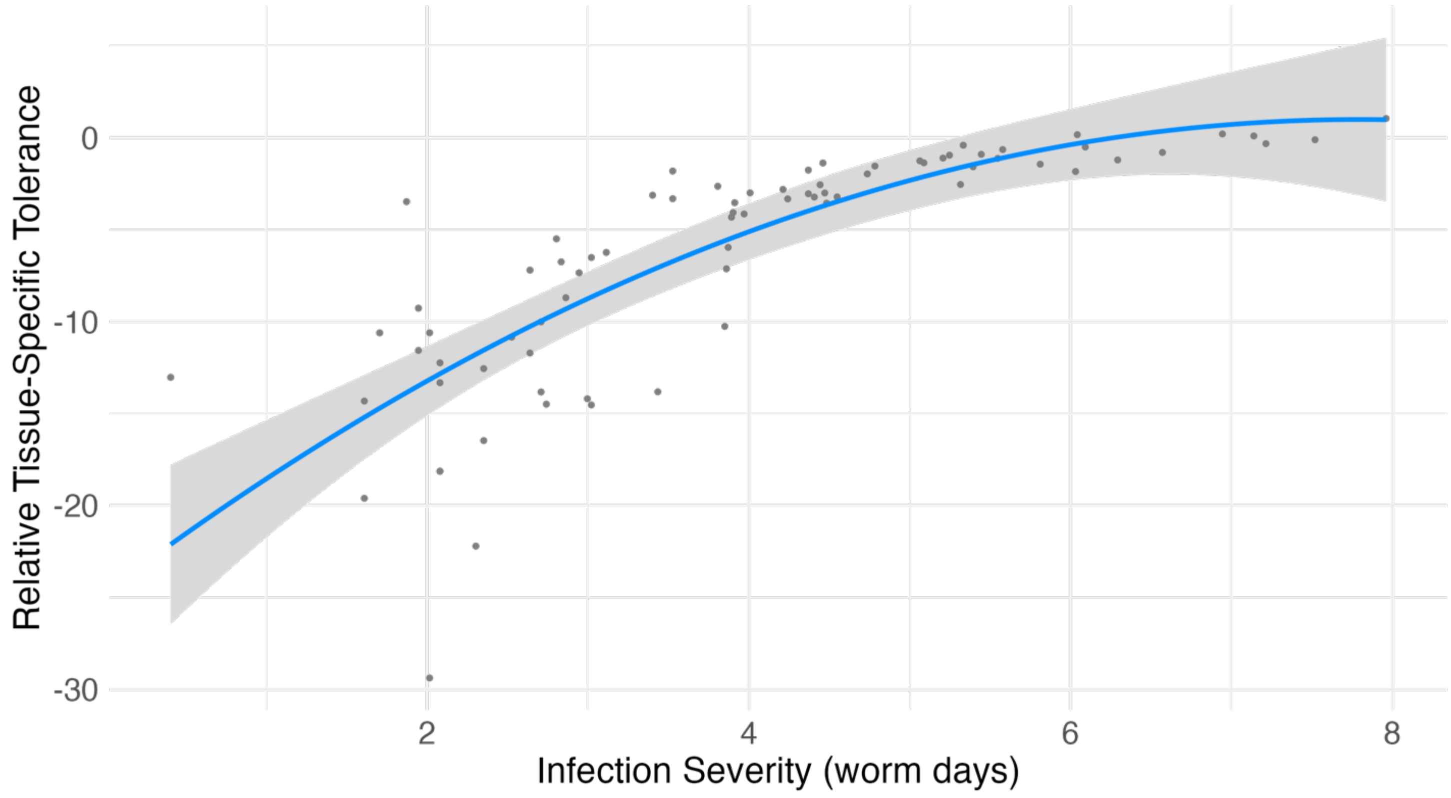

Figure S1. Polynomial regression demonstrates a trade-off between relative tissue-specific tolerance and parasite resistance (the inverse of infection severity). Points are back-transformed partial residuals from the model, and curves and shading give the model fits and 95% confidence bands.

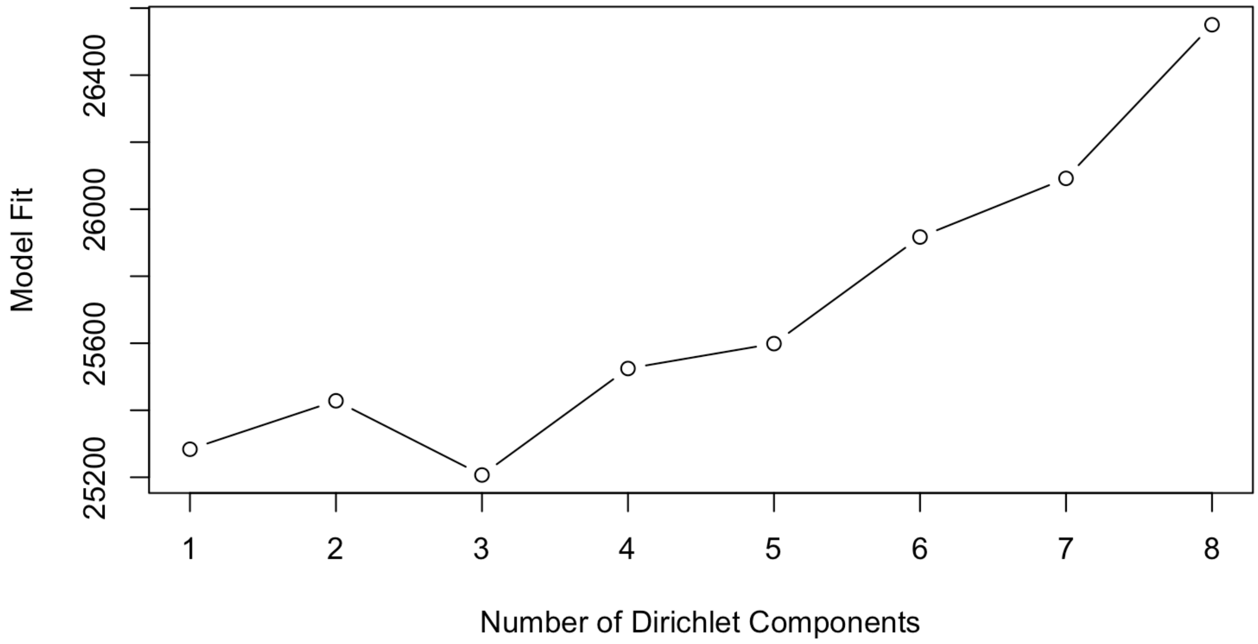

Figure S2. Identifying number of community types in pre-infection skin swabs of Guppies (n=50). Model fit (LaPlace approximation) with the number of Dirichlet components (k=3) was the lowest and best fit.

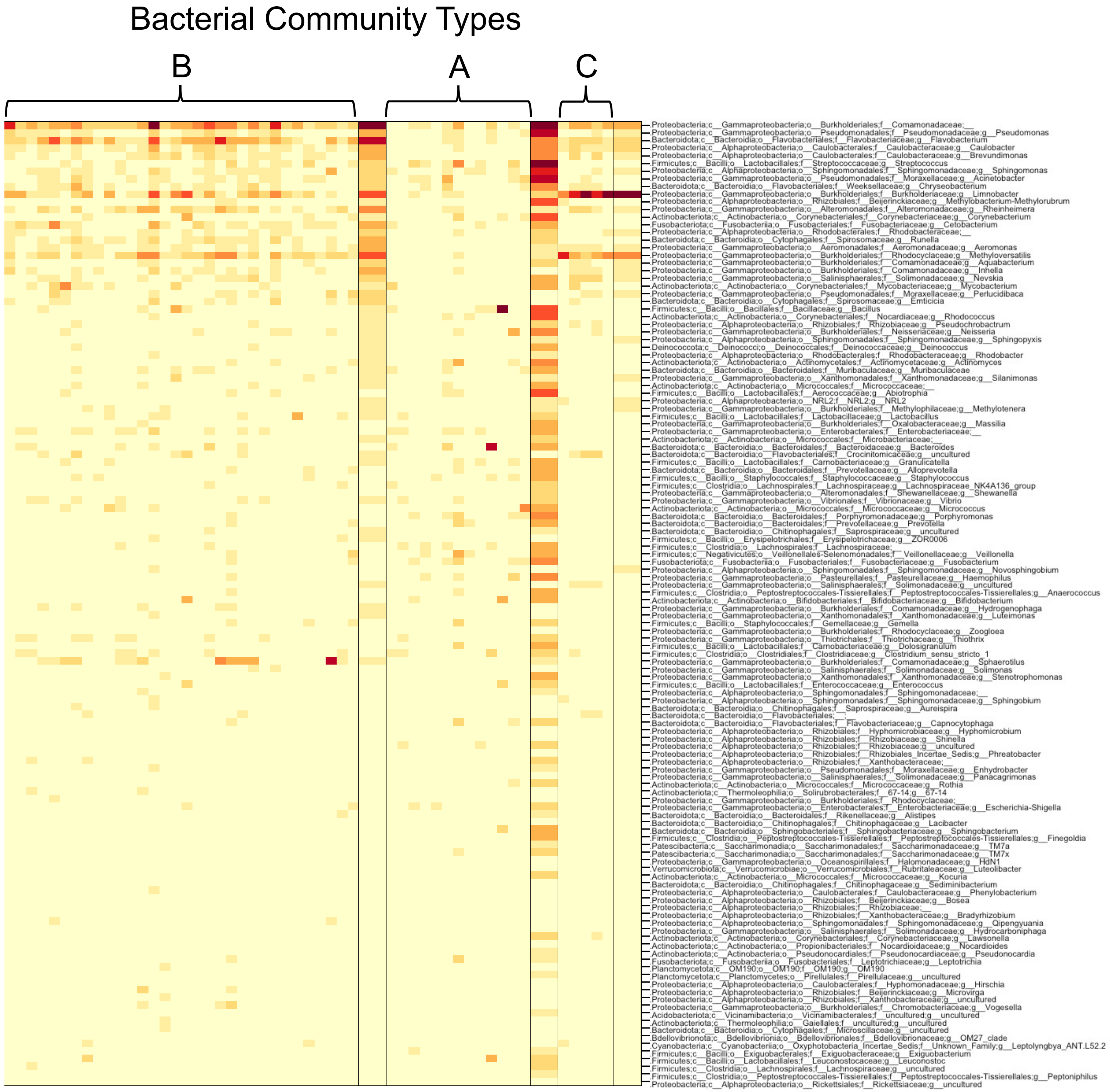

Figure S3. Top contributing Taxa the form the three Dirichlet components. Narrow columns are samples. Broader columns are the community types. Color represents square-root counts, with darker colors representing larger counts.

Table 1. Summary All Populations.

| Population | $N$ | $A_N$ | $A_R$[b] | $H_O$[c] | $H_S$[d] | $F_{IS}$ |
|---|---|---|---|---|---|---|
| Inf_Exp Control | 42 | 6.42 | 6.206 | 0.435 | 0.522 | 0.167 |
| Inf_Exp_Resistant | 37 | 7.143 | 7.102 | 0.496 | 0.512 | 0.034 |
| Inf_Expe_Susceptible | 36 | 4.28 | 4.27 | 0.432 | 0.484 | 0.108 |
| Wild Caught | 66 | 13.42 | 11.24 | 0.513 | 0.571 | 0.102 |

[a] Number of individuals genotyped (FSTAT)

[b] The number of alleles per locus (FSTAT)

[c] Allelic richness (FSTAT: based on minimum sample size of 59 individuals)

[d] Observed heterozygosity (GenAlEx)

[e] Gene diversity

Table 2. Estimates of Identity disequilibrium ($g_2$) for the four populations. P is the probability that the $g_2$ statistic is significantly greater than 0 when compared to a random distribution of genotypes. Calculated in Inbreed R (999 permutations).

| Population | $g_2 \pm$ SE | P |
|---|---|---|
| Wild Caught | -0.008 ± 0.01 | 0.691 |
| Control | 0.007 ± 0.035 | 0.463 |
| Resistant | 0.02 ± 0.036 | 0.2202 |
| Susceptible | 0.05 ± 0.05 | 0.08 |
