## Supplementary material for "Host genetics and the Skin Microbiome Independently Predict Parasite Resistance": Code Supplement

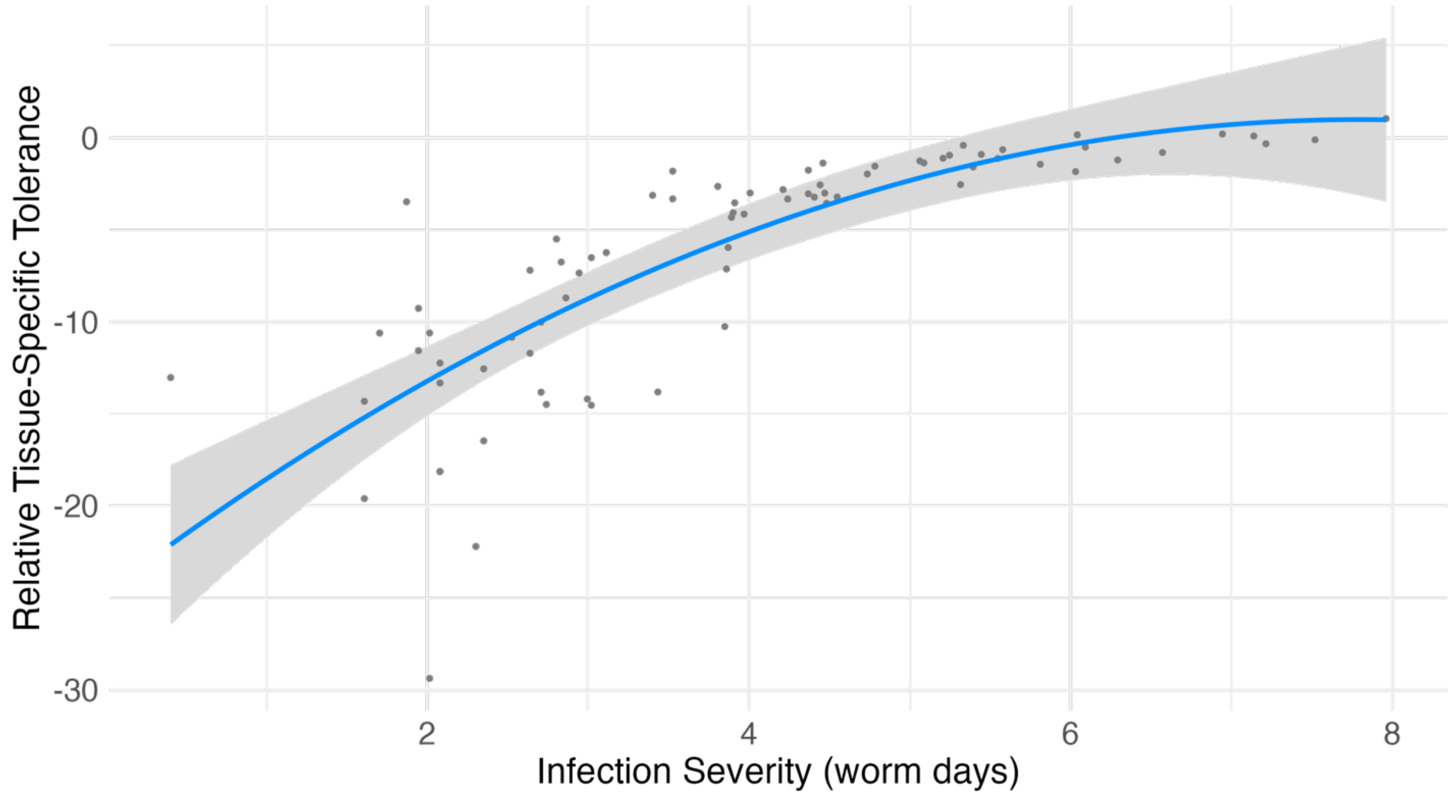


Figure S1. Polynomial regression demonstrates a trade-off between relative tissue-specific tolerance and parasite resistance (the inverse of infection severity). Points are back-transformed partial residuals from the model, and curves and shading give the model fits and 95% confidence bands.

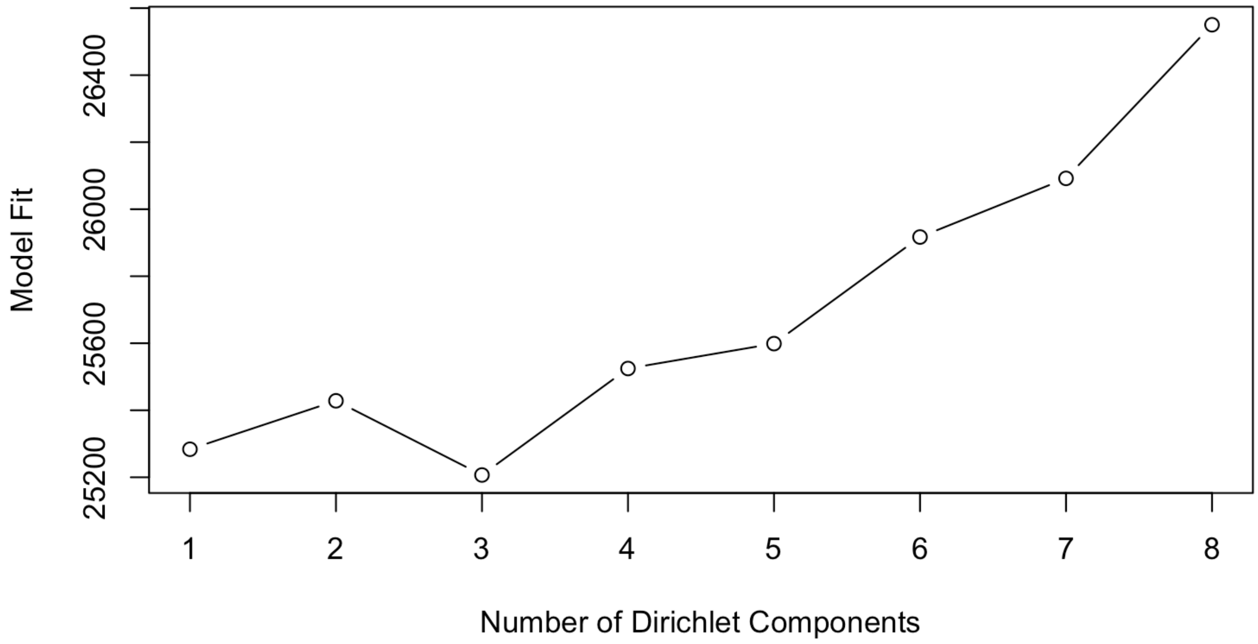


Figure S2. Identifying number of community types in pre-infection skin swabs of Guppies (n=50). Model fit (LaPlace approximation) with the number of Dirichlet components (k=3) was the lowest and best fit.

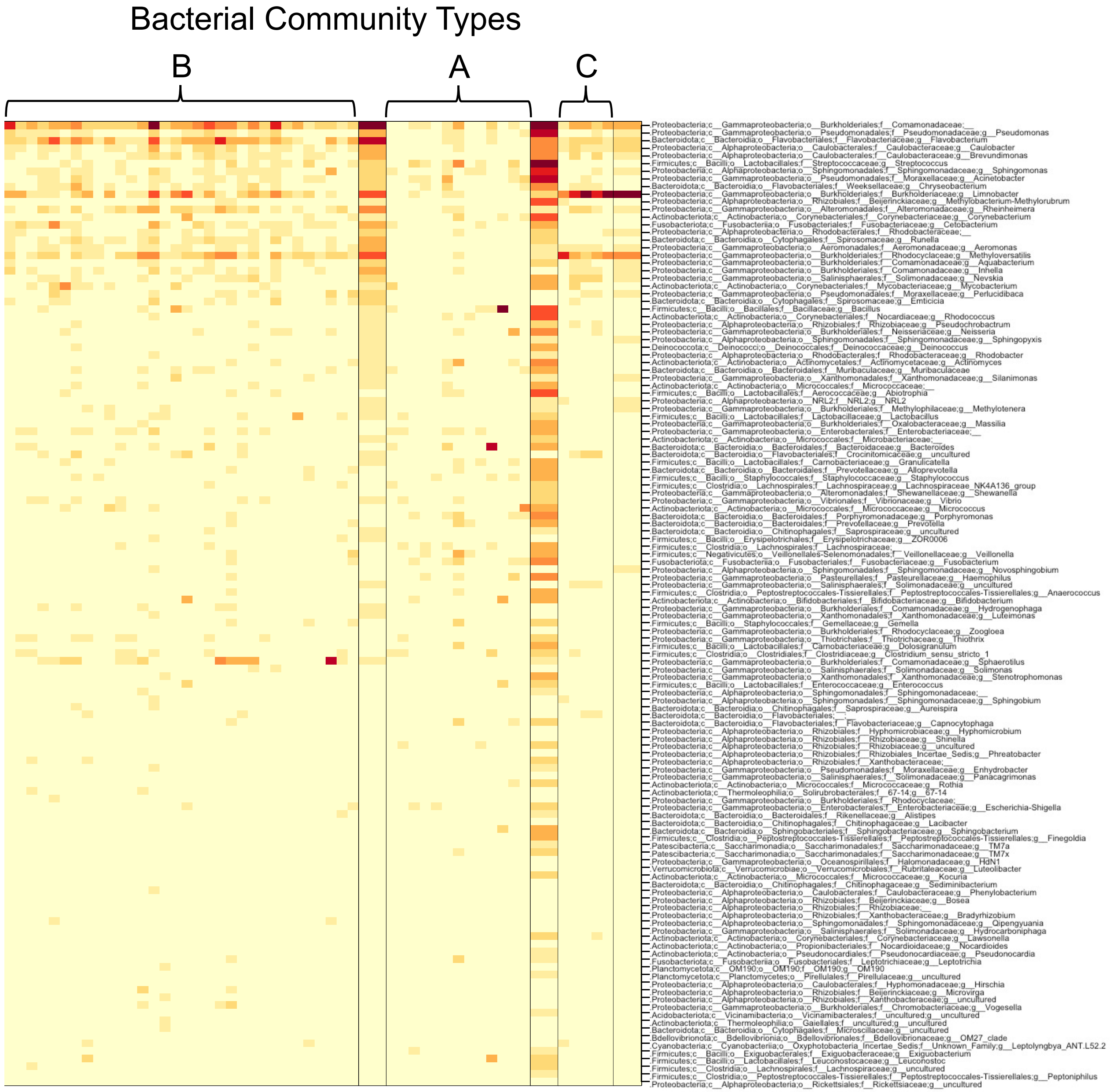


Figure S3. Top contributing Taxa the form the three Dirichlet components. Narrow columns are samples. Broader columns are the community types. Color represents square-root counts, with darker colors representing larger counts.

Table 1. Summary All Populations.

| Population | $N$ | $A_N$ | $A_R$[b] | $H_O$[c] | $H_S$[d] | $F_{IS}$ |
|---|---|---|---|---|---|---|
| Inf_Exp Control | 42 | 6.42 | 6.206 | 0.435 | 0.522 | 0.167 |
| Inf_Exp_Resistant | 37 | 7.143 | 7.102 | 0.496 | 0.512 | 0.034 |
| Inf_Expe_Susceptible | 36 | 4.28 | 4.27 | 0.432 | 0.484 | 0.108 |
| Wild Caught | 66 | 13.42 | 11.24 | 0.513 | 0.571 | 0.102 |

[a] Number of individuals genotyped (FSTAT)

[b] The number of alleles per locus (FSTAT)

[c] Allelic richness (FSTAT: based on minimum sample size of 59 individuals)

[d] Observed heterozygosity (GenAlEx)

[e] Gene diversity

Table 2. Estimates of Identity disequilibrium ($g_2$) for the four populations. P is the probability that the $g_2$ statistic is significantly greater than 0 when compared to a random distribution of genotypes. Calculated in Inbreed R (999 permutations).

| Population | $g_2$ ± SE | P |
|---|---|---|
| Wild Caught | -0.008 ± 0.01 | 0.691 |
| Control | 0.007 ± 0.035 | 0.463 |
| Resistant | 0.02 ± 0.036 | 0.2202 |
| Susceptible | 0.05 ± 0.05 | 0.08 |
